# High-throughput feedback-enabled optogenetic stimulation and spectroscopy in microwell plates

**DOI:** 10.1101/2022.07.13.499906

**Authors:** William Benman, Saachi Datta, David Gonzalez-Martinez, Gloria Lee, Juliette Hooper, Grace Qian, Gabrielle Leavitt, Lana Salloum, Gabrielle Ho, Sharvari Mhatre, Michael S. Magaraci, Michael Patterson, Sevile G. Mannickarottu, Saurabh Malani, Jose L. Avalos, Brian Y. Chow, Lukasz J. Bugaj

## Abstract

The ability to perform sophisticated, high-throughput optogenetic experiments has been greatly enhanced by recent open-source illumination devices that allow independent programming of light patterns in single wells of microwell plates. However, there is currently a lack of instrumentation to monitor such experiments in real time, necessitating repeated transfers of the samples to stand-alone analytical instruments, thus limiting the types of experiments that could be performed. Here we address this gap with the development of the optoPlateReader (oPR), an open-source, solid-state, compact device that allows automated optogenetic stimulation and spectroscopy in each well of a 96-well plate. The oPR integrates an optoPlate illumination module with a module called the optoReader, an array of 96 photodiodes and LEDs that allows 96 parallel light measurements. The oPR was optimized for stimulation with blue light and for measurements of optical density and fluorescence. After calibration of all device components, we used the oPR to measure growth and to induce and measure fluorescent protein expression in *E. coli*. We further demonstrated how the optical read/write capabilities of the oPR permit computer-in-the-loop feedback control, where the current state of the sample can be used to adjust the optical stimulation parameters of the sample according to pre-defined feedback algorithms. The oPR will thus help realize an untapped potential for optogenetic experiments by enabling automated reading, writing, and feedback in microwell plates through open-source hardware that is accessible, customizable, and inexpensive.

## Introduction

Optogenetic tools allow precise control of molecular activity inside cells using light as a stimulus. Because light can be readily interfaced with computers, optogenetic experiments are highly amenable to automation. Recently, due to the accessibility of small and programmable light sources, integrated circuits, and additive manufacturing, several groups have developed custom devices to miniaturize and parallelize optogenetic experiments^1–9^. These devices comprise arrays of light-emitting diodes (LEDs) positioned in the format of standard biological multi-well plates, often controllable by open-source hardware and software (e.g. Arduino, Python). Collectively, such devices allow programmable, high-throughput control of biological systems including in bacteria, yeast, mammalian cells, and other model organisms, with up to 3 stimulation colors per well. They are also inexpensive, often costing well below $1000 to construct. As a result, open-source devices for high-throughput optogenetics help realize the high potential of optogenetics for systematic, data-rich, and robust experiments without the need for complex robotics or bespoke microfluidics.

Despite the proliferation of such hardware, there is a lack of devices that also allow simultaneous measurement of the samples, for example through spectroscopy. Simultaneous stimulation and measurement in one integrated device would be highly enabling. First, it would streamline optogenetic experiments, removing the need for manually transferring the sample plate from the stimulation device to a plate reader or microscope, allowing higher resolution time sampling and removing sources of experimental error, such as unwanted light exposure while transferring samples. Second, real-time measurement and perturbation would allow computer-in-the-loop feedback control, where the stimulation can be adjusted based on the current state of the sample. Such control could have many uses, for example for optimizing optogenetically controlled metabolic pathways in which enzymes are expressed at precise levels and only during a particular phase of cell culture growth^10^. Although all-optical feedback control has been previously implemented, it has required expensive microscopes^11, 12^ or customization of flow cytometers^13, 14^ and could only act on one sample at a time. Recently, custom devices were reported that allowed optogenetic stimulation and imaging of bacterial cultures in microwell plates^15^ or in batch culture^16^. However, these devices were limited to experiments in four wells or one bulk culture, respectively.

In recent years, open-source spectrophotometers have been described that could in principle be coupled with illumination devices. Richter et al demonstrated that a Tecan plate reader could be retrofitted for optogenetic stimulation by converting the on-board fluidics machinery to position an LED-coupled optical fiber above predefined wells^17^. However, this approach could only read and write from one sample at a time and required access to a Tecan plate reader that could be customized. Separately, Szymula et al described the open-source plate reader (OSP), which provides full-spectrum absorbance and fluorescence detection in microwell plates and allows optogenetic stimulation and reading of an individual well^18^. However, this instrument could not regulate sample temperature and thus could not support continuous cell culture, and also required sequential stimulation/measurement of each well. Jensen et al developed a 96-well solid-state plate-reader that used an array of 96 phototransistors to optically measure each well independently^19^. This device could measure light from all wells simultaneously and could be shaken and multiplexed within bacterial incubators. However, it could only measure OD but not fluorescence, and could not be used to stimulate optogenetic systems.

In this work, we describe the development of the optoPlateReader (oPR), an integrated device that allows 96 parallel channels of optical stimulation, measurement of fluorescence and optical density, and feedback control of stimulation based on real-time measurements of biological samples. We characterize the detection limits of our device and demonstrate its ability to measure bacterial growth and arabinose- or light-inducible expression of the fluorescent protein mAmetrine with low variability between wells. Finally, we demonstrate that 96 separate cultures can be independently and simultaneously regulated to control gene expression programs in real time conditional on the current growth or expression state of the sample.

## Results

### Design of optoPlateReader for simultaneous optogenetic reading and writing

The optoPlateReader (oPR) was designed for high-throughput light stimulation with real-time fluorescence and absorbance measurements in a 96-well plate format. Other important specifications included 1) a small footprint such that the device could be placed into a standard cell culture incubator for environmental control, 2) stable mechanical and electrical connections for robustness and to allow shaking, 3) integration between measurement and stimulation to allow for autonomous feedback control, 4) a user interface that allowed easy programming of all experimental parameters.

The oPR is composed of two separate device modules: the optoPlate, which provides light sources for optogenetics and OD readings, and the optoReader, which provides components for optical measurement and light sources for fluorescence excitation (**Figure 1A**). All stimulation and measurement can be controlled independently for each of the 96 separate wells. Both device modules consist of a custom-designed printed circuit board (PCB) assembled with surface-mounted semiconductor components. Surface-mounted components can be small in size, allow for rapid and precise device assembly without the need for specialized equipment or expertise in hand soldering (See **Methods** and **design files),** and offer the potential for scalable production. A clear-bottom, opaque-walled 96-well sample plate is positioned between the optoPlate (top) and the optoReader (bottom) modules. Both modules are fitted with 3D-printed adapters that securely mate the circuit boards to the sample plate. The small format of the assembled oPR allows it to fit within cell culture incubators, and the lack of moving parts and wires provides robustness, for example allowing the device to be shaken. For shaking, the oPR can be mounted on a microplate orbital shaker. The oPR can communicate with the shaker via a 5V relay, allowing shaking to be paused during measurements and resumed after measurements are complete. For environmental control, the device assembly can be placed inside of a standard 37°C cell culture incubator.

**Figure 1.**
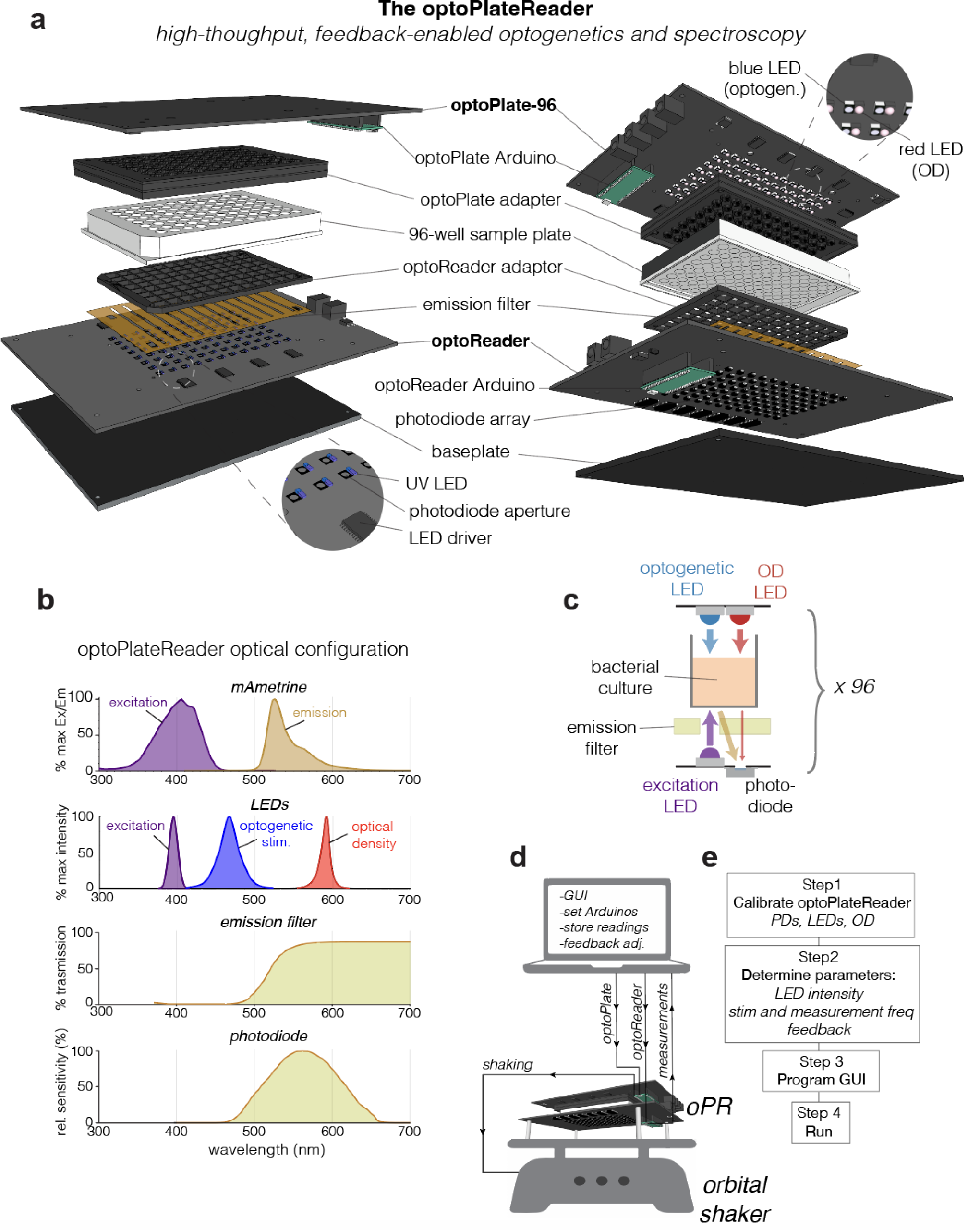
The optoPlateReader (oPR) enables high throughput optical stimulation and spectroscopy in a 96-well format. **a** Exploded view of the oPR. The optoPlate-96 and optoReader each feature an array of 96 LEDs or photodiodes that independently optically stimulate and measure each well of a 96-well plate. The 3D-printed adaptors provide light insulation between wells and provide a tight mating to a black-walled, clear-bottom 96-well plate. **b** Optical spectra for mAmetrine and the oPR components, which were selected to maximize detection of fluorescence emission and absorbance measurements while preventing detection of excitation light. **c** Schematic of optical stimulation and measurement of a single well in the oPR. **d** Communication of a computer with the oPR during an experiment. The computer receives user input and real-time measurements from the oPR, calculates updates to stimulation parameters based on feedback algorithms, and sends commands and updated protocols to the optoPlate and optoReader Arduinos. The optoReader Arduino sends photodiode measurements back to the computer. Communication between the oPR and an orbital shaker allows for sample agitation between measurements. **e** Overview of workflow for oPR demonstrating steps a user must take after oPR construction and before performing an experiment. Note that whereas calibration of blue LEDs, UV LEDs, and photodiodes needs to be performed once, calibration of OD measurements (red LEDs) must be performed before each experiment.

The optoPlate is derived from a previously reported optogenetic stimulation device, the optoPlate-96^1^. The optoPlate consists of 96 pairs of LEDs, arranged such that each pair of LEDs illuminates a single well of a 96-well plate. We adapted this device to rest on top of the sample well plate, and we selected LED pairs to allow both optogenetic stimulation (470 nm) and optical density readings (600 nm). We selected bi-color blue/red LEDs for optogenetic stimulation because the majority of current optogenetic tools respond to blue or red light, though we exclusively used blue light for optogenetic stimulation in this study. The bi-color LED can be replaced with a mono- or bi-color LED of any wavelength for custom applications, provided that the LED form factor is compatible with the optoPlate PCB (either PLCC2 or PLCC4). A 3D-printed adapter is mounted on the optoPlate to securely mate and align the optoPlate with the sample plate. The intensity of all LEDs is regulated via pulse width modulation (PWM) enabling 4095 levels of intensity control.

The optoReader is a solid state, 96-well fluorescence plate reader (**Figure 1A)**. The optoReader contains 96 pairs of one photodiode and one UV LED, arrayed in 96-well format. The photodiodes are the light-sensing element used to measure fluorescence and OD. The optical configuration of the oPR has been optimized to measure fluorescence of mAmetrine. mAmetrine is a derivative of GFP with a long Stokes shift^20^— that is, with a relatively large difference between its excitation and emission wavelength (**Figure 1B, top**). The long Stokes shift allows the excitation light (UV) to be efficiently filtered out while maximally preserving the emission light using only inexpensive filters (see below and **Methods**). An additional benefit of mAmetrine is that its excitation spectrum minimally overlaps with the blue LED spectrum, minimizing bleaching of the fluorescent protein from optogenetic stimulation (**Figure 1B**). To excite mAmetrine, we used a near-UV LED (395 nm) positioned next to each photodiode. To filter out UV excitation light from the photodiode detector, we implemented emission filters above all photodiodes using canary yellow camera filters (Rosco Roscolux) (**Figure 1A,B**). We cut apertures in these filter films above the UV LEDs such that the UV light could be transmitted onto the sample, but its reflection onto the photodiodes would be attenuated (**Figure 1C**). For further filtering, we selected photodiodes that had minimal responsivity to light below 450 nm, further attenuating signal from the UV LED while permitting detection of light from mAmetrine emission or the OD LED (**Figure 1B)**. For further possible modification, the optoReader can also accommodate an additional LED component in each well position, if desired, for example for detection of fluorescence from multiple fluorophores. As with the optoPlate, a 3D-printed adapter mates the optoReader and the bottom of a 96-well plate, providing stability, light insulation between wells, and alignment between all modules of the assembled oPR. Lack of light leakage between wells, as well as spatial homogeneity of illumination, were experimentally confirmed (**Supplementary Figure 1**)

Both the optoPlate and optoReader modules are driven by on-board Arduino microcontrollers that communicate with the local LEDs and photodiodes, with the central computer, and with the shaker. LEDs are controlled by serial communication through 24-channel LED driver chips, as used previously^2^. To read 96 analog signals from all photodiodes, six 16-channel multiplexers take sequential readings from individually addressable wells, and readings are transmitted to one analog input pin on the Arduino. For more details on the optoReader circuitry, see **Supplementary Figure 2**. Both Arduinos communicate with a central computer to send and receive commands through USB communication. The computer runs custom Python software that sends illumination/measurement parameters to the Arduinos, coordinates timing (e.g. to ensure that optogenetic stimulation does not occur during readings), and stores and processes measurements (**Figure 1D**). The oPR is able to record 96 fluorescence or OD readings in <1 min. The ability for the oPR to rapidly measure a sample, perform calculations on those values, and dynamically update stimulation parameters enables computer-in-the-loop feedback control, where optogenetic stimulation can be modified in real time based on the current state of the sample. Such feedback control can be implemented in 96 independent experiments at the same time.

We provide all design files and a parts list to print and assemble a fully functional oPR (See **Methods**). With all components in hand, a fully-functional oPR can be assembled in ∼6 hours for ∼$700, with price decreasing if components are purchased in larger quantities. After assembly, the general steps to perform an experiment are as follows (**Figure 1E**): First, all oPR components must be calibrated to allow measurement and stimulation with minimal variation between wells (see below). Second, the experimental cells must be grown and plated, and the experimental conditions (stimulation intensity and frequency, measurement frequency) must be determined. Third, the full oPR device with sample plate must be assembled and powered, the OD readings must be calibrated, and the experimental parameters must be entered into the graphical user interface (GUI). Finally, the experiment is initiated from the GUI. The following sections detail the oPR software and calibration protocols and provide examples for the types of experiments that can be performed with the oPR.

### The oPR software

The oPR software allows the user to define all experimental parameters, coordinates the timing of all electronic components, takes and stores measurements, and runs feedback algorithms to adjust stimulation parameters based on predefined specifications (**Figure 2A**). The GUI allows for easy programming of all stimulation, measurement, and feedback parameters within each of the individual 96 wells (**Figure 2B**). The GUI home screen features three functions: 1) “Calibrate OD”, 2) “Calibrate Blue”, and 3) “Start Experiment”. The calibration buttons allow for automated calibration of the oPR components (see next section). After calibration, the “Start Experiment” button leads to a window titled “Stimulation Protocol”, which prompts the user to define optogenetic stimulation protocols and to assign those protocols to individual wells or groups of wells. For each protocol, the user can specify the intensity of the blue LED, the duration that the LED will be ON, and the subsequent duration that LED will be OFF. The LED will loop through these ON and OFF durations continuously. Up to 96 distinct protocols can be specified. The user can also specify a feedback function to be applied to each pattern of wells. Arbitrary feedback inputs, outputs, and algorithms can be programmed in the FeedbackFuncs.py file (see **design files** and **manual**). In the subsequent window, the user defines measurement parameters for the optoReader, specifically the duration and frequency of OD and fluorescence readings. The same measurement parameters are applied to all wells. For each type of measurement, the user can specify the number of individual readings that will be averaged in order to minimize measurement noise. For the studies in this report, we averaged 100 readings per measurement. In the final window, the GUI allows the user to review and edit the experimental protocols before running the experiment. At the start of the experiment, the software generates .csv files in which the OD and fluorescence measurements will be recorded and updated. At the end of the experiment, the user retrieves these .csv files and processes them as needed for data analysis and visualization.

**Figure 2.**
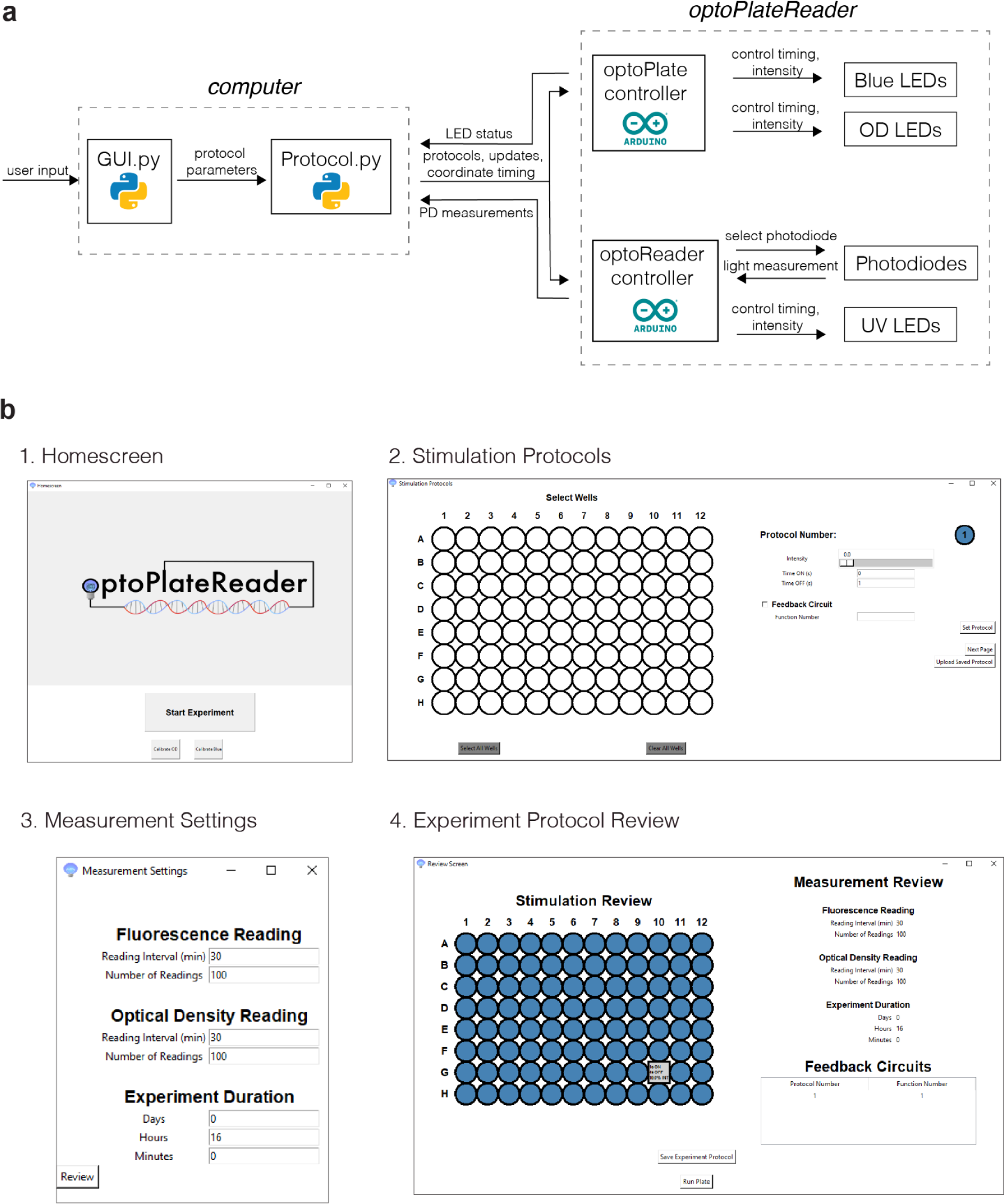
oPR software workflow and graphical user interface (GUI) for programming oPR protocols. **a** Software architecture for communication between user, computer, and oPR. The GUI and Protocol python scripts send user-defined stimulation and measurement protocols to the optoPlate and optoReader Arduinos that control individual components in both device modules. Photodiode measurements collected by the optoReader Arduino are returned to the computer for reporting or feedback-driven signal adjustments. **b** Windows of the GUI. (1) The user can automatically calibrate OD and Blue LEDs from the homescreen. (2) Wells, light dose, illumination timing, and feedback algorithm are specified for up to 96 independent protocols. (3) Photodiode reading frequency parameters are specified. (4) The user can review and save their experimental protocols prior to running the experiment.

### Calibrating the optoPlateReader

Each of the four optical elements of the oPR (UV LED, blue LED, OD LED, photodiode) must be calibrated to minimize measurement noise that originates from variability during component manufacture or device assembly (**Figure 3**). Each set of components can be calibrated in an automated manner using the GUI (OD and Blue LEDs) or in a semi-automated manner using files available in the **oPR Repository** (See **Methods**) (UV LEDs and Photodiodes). Calibration involves measuring each of the 96 components, calculating their variability (coefficient of variation (CV), the standard deviation normalized by the mean), calculating normalization factors to minimize CV, performing new measurements with the normalization factors, and iterating over multiple rounds until CV is minimized. We reasoned that, because each well contains a light detector (the photodiode), we could first calibrate the photodiodes to an external, uniform light source and then subsequently calibrate each LED using the calibrated photodiodes. A detailed description of all calibration procedures can be found in the **Methods** section.

**Figure 3.**
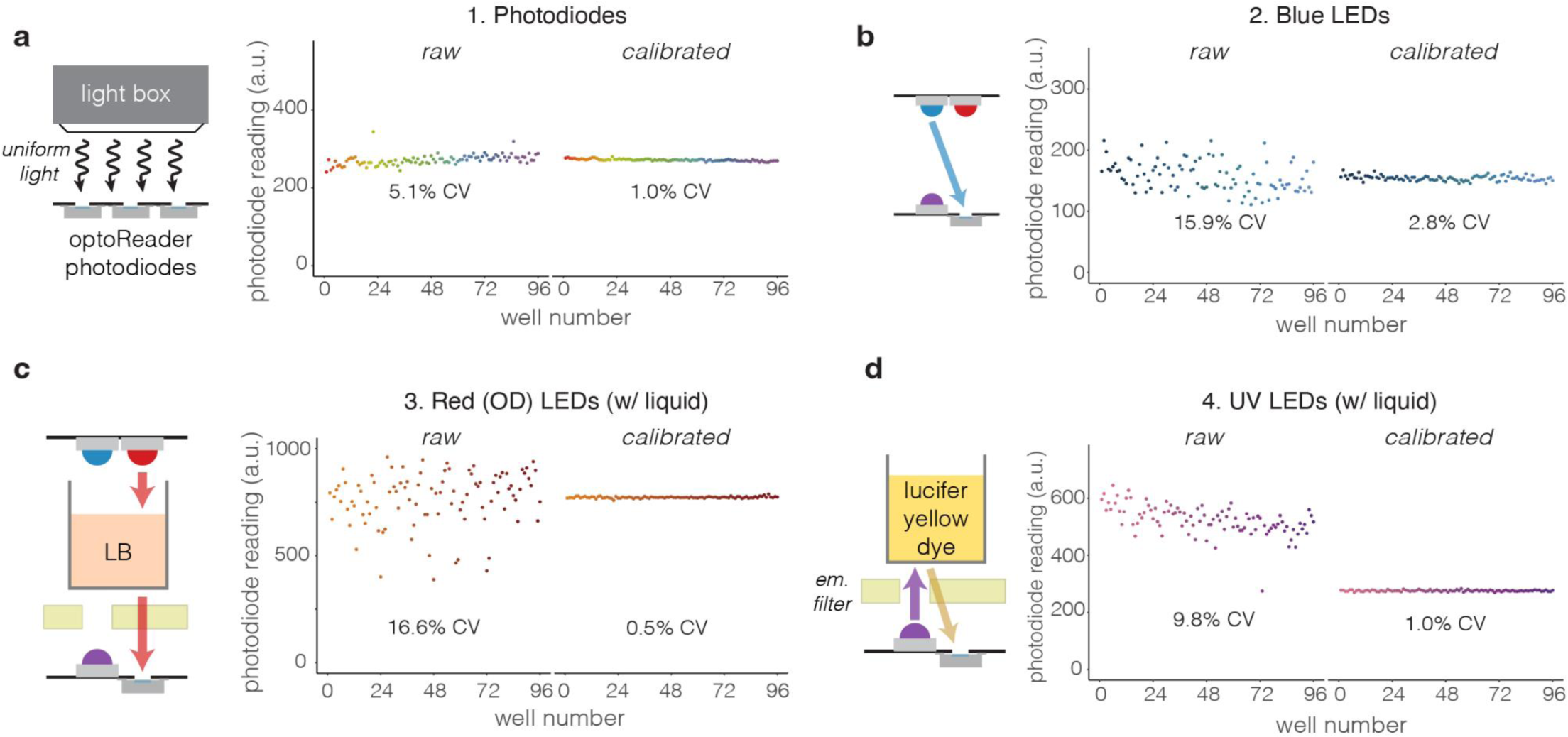
Component calibration for precise stimulation and measurement between wells. **a** Calibration of photodiodes must be performed first in order to use photodiodes to calibrate the LED components. Photodiodes are calibrated by using a uniform white light source to illuminate all wells with the same light intensity. **b** Blue LEDs are calibrated with the photodiodes in a fully assembled oPR, using an empty 96-well plate and no emission filter. **c** OD LEDs are calibrated in a fully-assembled oPR with an equal volume of LB in each well to to account for light refraction through the medium. This calibration should be performed at the start of each experiment using the “Calibrate OD” function found in the GUI. **d** UV LED calibration is performed in a fully-assembled oPR with an equal volume of diluted lucifer yellow dye in each well (40 µg/mL shown here). Calibration is performed by adjusting LED intensity to normalize variation in the measured emission intensity of the dye. In all cases, calibration reduced variation between wells to CV = 1-3%

To calibrate the photodiodes, we used an instrument that projected white LED light uniformly over a surface, and we further used diffuser film to homogenize illumination and adjust irradiance intensity, which we confirmed with a handheld power meter (see **Methods**) (**Figure 3A**). Uncalibrated optoReader photodiodes initially recorded 96 values with CV = 5.1%. After one round of calibration, we measured dramatically less variation between photodiodes (CV = 1.0%). Further calibration rounds yielded no further decrease in CV. Calibration factors were also found to be independent of light exposure intensity (**Supplementary Figure 3**).

We then used the calibrated photodiodes to calibrate the blue, OD, and UV LEDs in the fully assembled oPR. Blue LED variability (CV = 15.9%) was reduced to 2.8% after two rounds of calibration (**Figure 3B**). The OD LEDs were calibrated in a similar manner except that LB was added to the wells to reproduce the light refraction caused by liquid in the wells, which we hypothesize could contribute to well-to-well variation in OD readings. After 3 rounds of calibration, OD LED variability was reduced from CV = 16.6% to 0.5% (**Figure 3C**). Finally, we calibrated the UV LEDs by measuring variability in fluorescence emission of Lucifer Yellow dye that was added to each well of a sample plate. Lucifer Yellow dye has similar fluorescence spectra to mAmetrine and thus is compatible with the optical configuration of the oPR. After 2 rounds of calibration, measurement variability decreased from CV = 9.8% to 1.0% (**Figure 3D**).

We note that the OD LED calibration should be performed at the beginning of each experiment since we have found significant variability in OD readings between experiments, likely due to slightly different sample refractive properties and device alignments. OD LED calibration is performed using the “Calibrate OD” function on the opening window of the GUI (**Figure 2B**). By contrast, calibration of the LEDs can be performed much less frequently, primarily to account for changes in LED brightness due to extended use. Calibration factors were robust to changes in temperature, allowing accurate readings to be taken at 37°C based on calibration factors obtained at room temperature (**Supplementary Figure 4**).

### Characterizing oPR measurements

With a fully calibrated oPR, we sought to characterize the limits and sensitivity of fluorescence signal detection. We generated a dilution series of Lucifer Yellow dye (2-250 µg/mL), and we measured each concentration in every well of a 96-well plate (**Figure 4, Supplementary Figure 5**). Fluorescence readings were highly consistent between the 96 wells. Readings increased monotonically with concentration, with a linear region extending to 40 µg/mL (**Figure 4A**). The average lower limit of detection (LOD) (1.0 +/- 0.2 µg/mL) was also highly consistent between wells (**Figure 4B**), yielding an average dynamic range of 40 (**Figure 4B**). Measurement sensitivity — or, the relationship between the photodiode counts and concentration — showed little variation between wells (12.5 +/- 0.3 counts/µg/mL, **Figure 4C**) and was highly linear (R^2^ = 0.996 +/- 0.002, **Figure 4D**). Collectively, these results demonstrate that the oPR can measure fluorescence in 96 wells simultaneously with high sensitivity, high precision, and low variance between wells. Of note, although the oPR was not as sensitive for fluorescence measurements as could be obtained by a commercial plate reader (LOD of Tecan Infinite M200 Pro: 12 +/- 0.3 ng/mL (**Supplementary Figure 6**)), the oPR was sufficiently sensitive for robust fluorescence measurements from bacterial cultures (see **Figures 6-8**).

**Figure 4.**
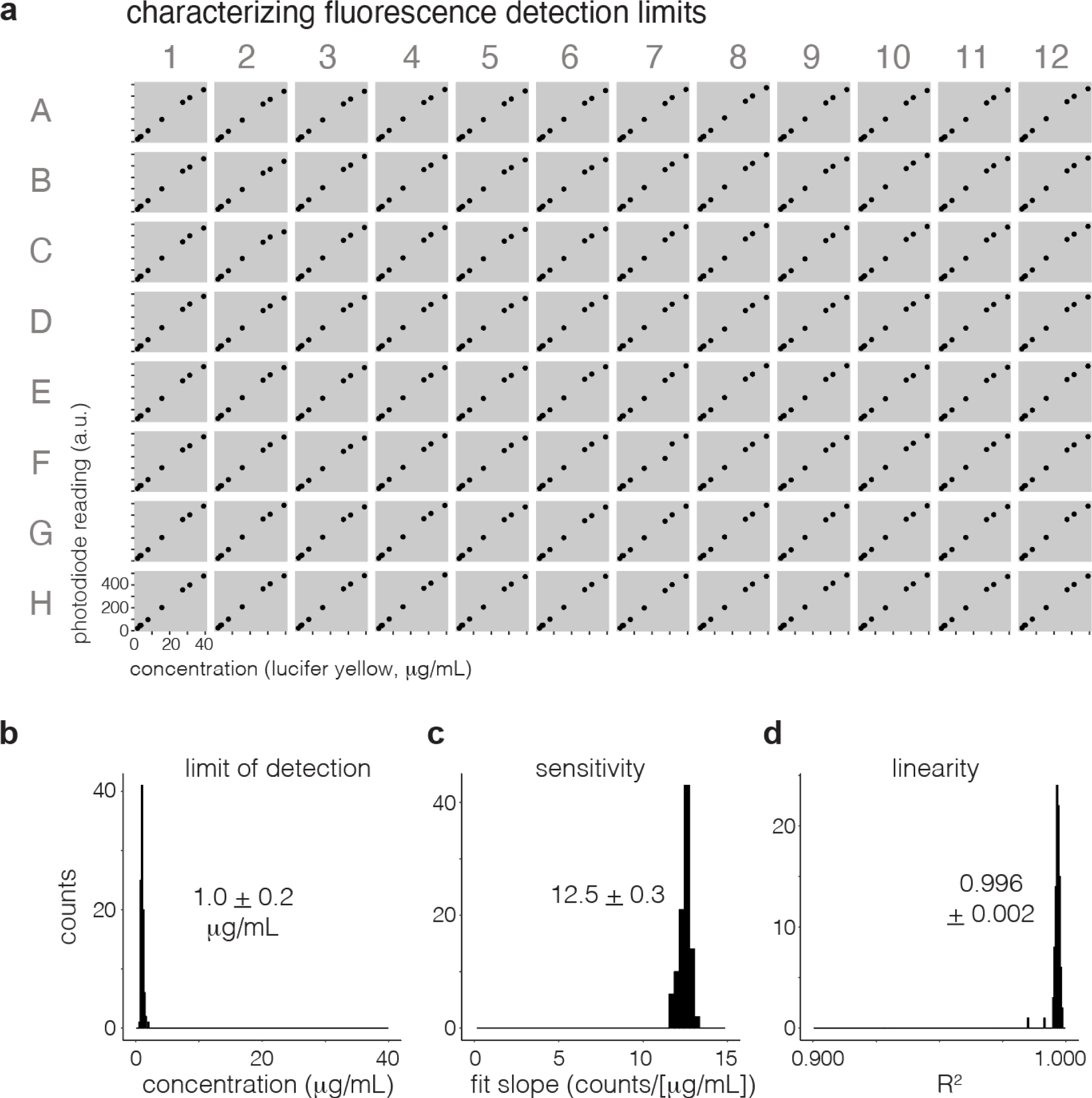
Characterization of fluorescence detection. **a** Fluorescence measurements of lucifer yellow dye were taken over a range of concentrations in each of 96 wells in a calibrated oPR. The linear range of measurements is shown (full concentration range in **Supplementary Figure 5**). **b** The average limit of detection of each well was 1.0 +/- 0.2 µg/mL, showing low variability between each of the 96 wells. **c** The sensitivity —or, the slope of the linear fit of the relationship between photodiode counts and concentration — also showed low variability between wells. **d** R^2^ values of the fits described in **(c)** confirm linearity in all wells. We next characterized the optical density measurement capabilities of the oPR by comparing oPR measurements to those obtained with a Tecan Infinite M200 Pro. We generated a twenty-fold dilution series of 1.1 µm polystyrene beads, a common reagent for OD measurement calibrations^21^. We plated each dilution into all wells of a 96-well plate and measured optical density in both the oPR and the Tecan in rapid succession. OD measurements from the oPR vs the Tecan measurements showed a highly linear relationship in all wells (R^2^ = 0.98 ± 0.004, Figure 5A-D. For details on OD calculation, see **Methods**).

**Figure 5.**
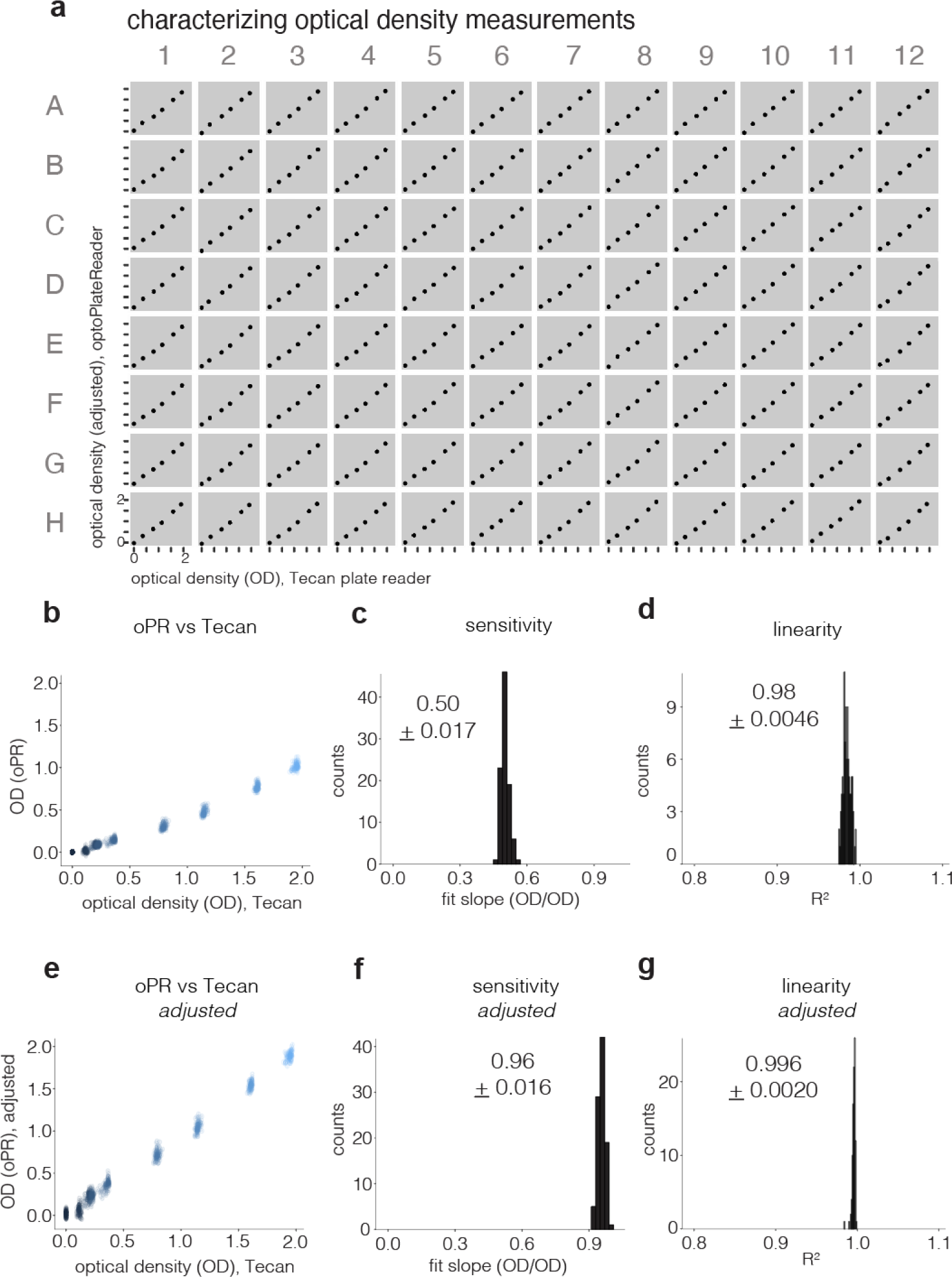
Characterization of optical density measurements. **a** oPR OD measurements were characterized by making dilutions of 1.1 µm diameter plastic beads, plating these dilutions in each well of a 96-well plate, and measuring OD on the oPR and on a Tecan Infinite M200 in rapid succession. Each Tecan reading is the average of 25 readings from the center of each well. Data shows calibrated/adjusted results, as described in **Methods**. **b** Raw data from (**a**) is overlaid. Each color represents a distinct dilution of beads. Each cloud has 96 data points representing each of 96 wells. **c** A histogram of the slope of linear fits of each plot in part (**a**). Average sensitivity is mean slope +/- 1 s.d. of 96 fits. **d** A histogram of R^2^ values of linear fits from (**b**). Average R^2^ is mean +/- 1 s.d. of 96 fits. **e** To allow direct comparison between Tecan readings and oPR, a transform function was calculated from data shown in **b-d** for each well. This transform was then applied to a second independent data set to generate “adjusted oPR readings” that closely matched Tecan readings. **f** Slopes of these adjusted data show high correspondence (slope ∼1) and high linearity (**g**) between oPR and Tecan OD readings.

**Figure 6.**
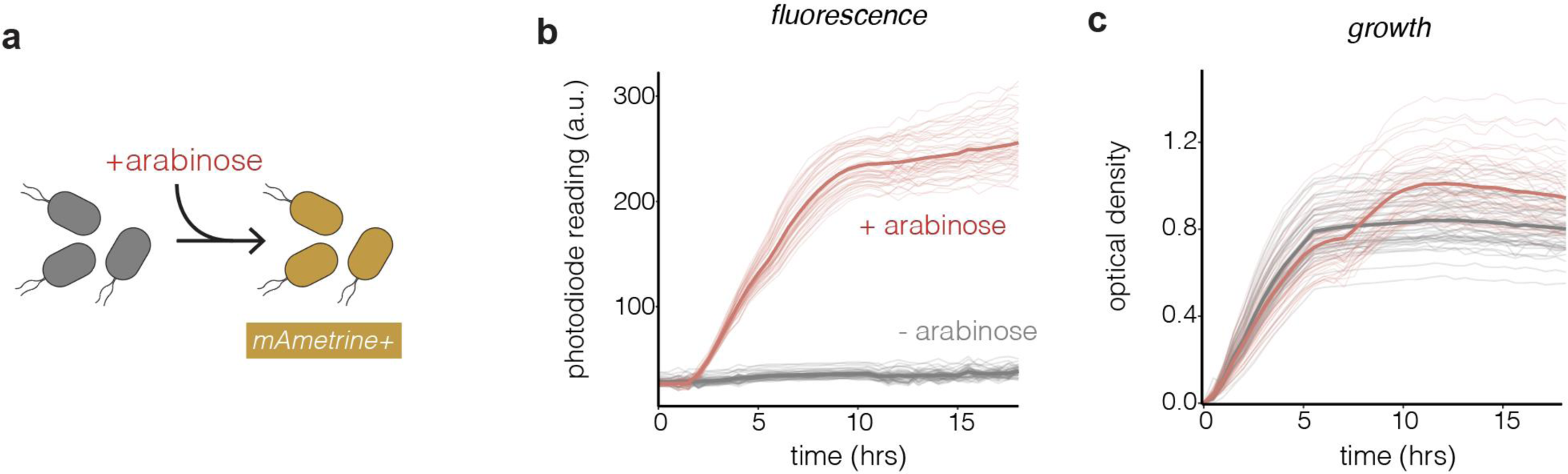
The oPR detects fluorescence and bacterial growth in real time. **a** The oPR was used to measure growth and fluorescence in *E. coli* that expressed mAmetrine under an arabinose-inducible promoter. **b** mAmetrine fluorescence was detected in every well that was treated with arabinose (red), and was not detected in the absence of arabinose (grey). Each trace corresponds to readings from an individual well. Bold traces represent means of all 48 replicates per condition. Fluorescence was measured every 20 min for 18 hrs. **c** OD readings of the same wells depicted in (**b**). Both measurements were taken every 20 min at 37 °C. Each trace corresponds to readings from an individual well. Bold traces represent means of all 48 replicates per condition. The increase in OD at ∼7 hours in the “+arabinose” wells is likely due to the bacteria switching to arabinose as a carbon source once glucose is exhausted^23^.

**Figure 7.**
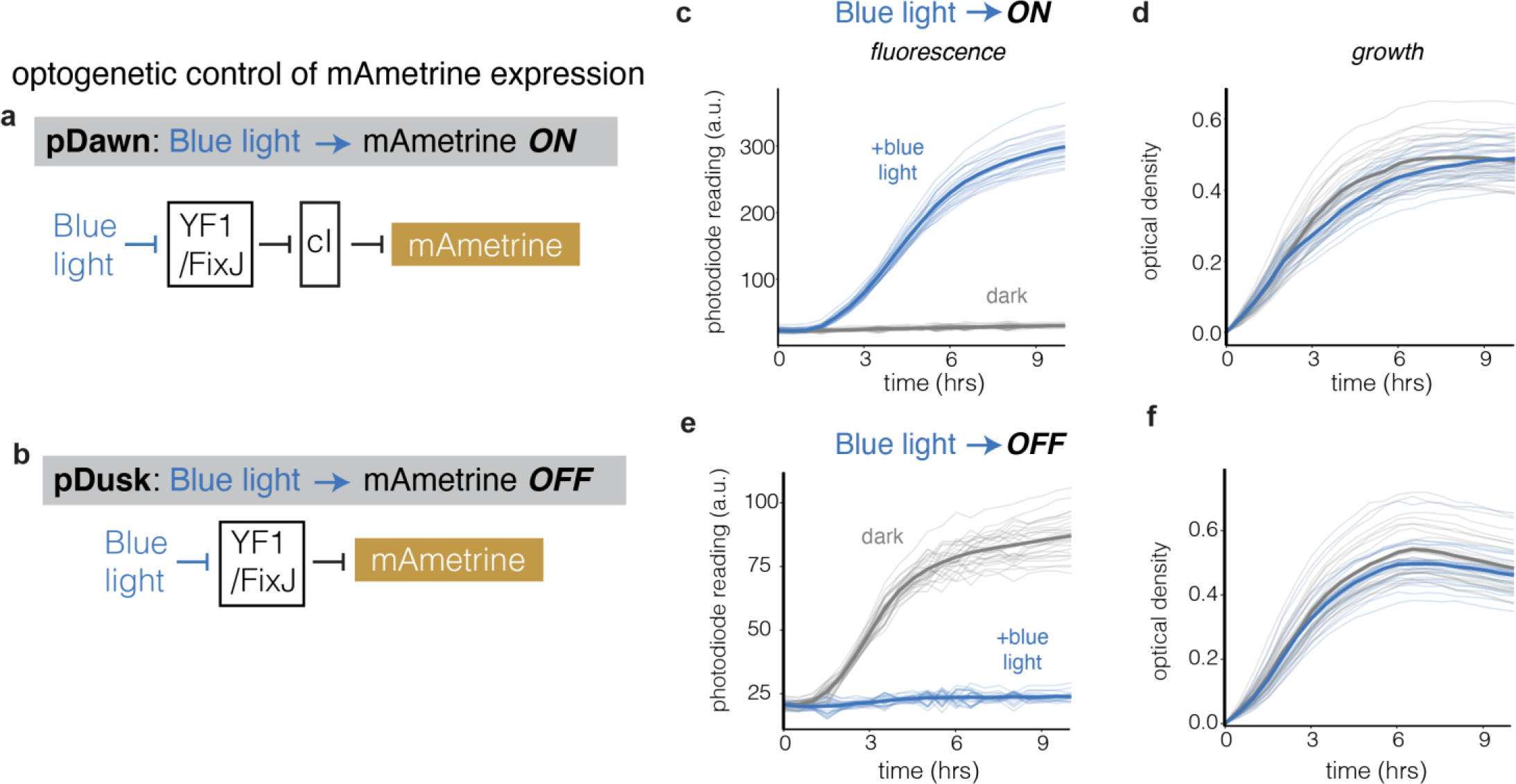
Simultaneous optogenetic stimulation and spectroscopy in growing bacteria. The oPR was used to stimulate and measure *E. coli* that expressed mAmetrine controlled by one of two optogenetically-controlled transcriptional circuits, pDawn (**a**) and pDusk (**b)**. In pDawn, blue light triggers mAmetrine expression, while in pDusk, blue light suppresses its constitutive expression. Fluorescence (**c**) and OD (**d**) of pDawn samples in response to blue light (blue) and dark (grey) conditions. The oPR detects mAmetrine only in samples exposed to blue light, demonstrating successful optogenetic induction. Traces represent measurements from individual wells, and bold traces represent means from each condition (24 replicates with blue light, 24 replicates in the dark). Data was collected every 30 minutes for 10 hrs at 37 °C. Fluorescence (**e**) and OD (**f**) of pDusk samples in response to blue light (blue) and dark (grey) conditions. The oPR detects mAmetrine only in samples that were not exposed to blue light, demonstrating successful optogenetic suppression. Traces represent measurements from individual wells, and bold traces represent means from each condition (24 replicates in the dark, 24 replicates with blue light). Data was collected every 30 minutes for 10 hrs at 37 °C.

**Figure 8.**
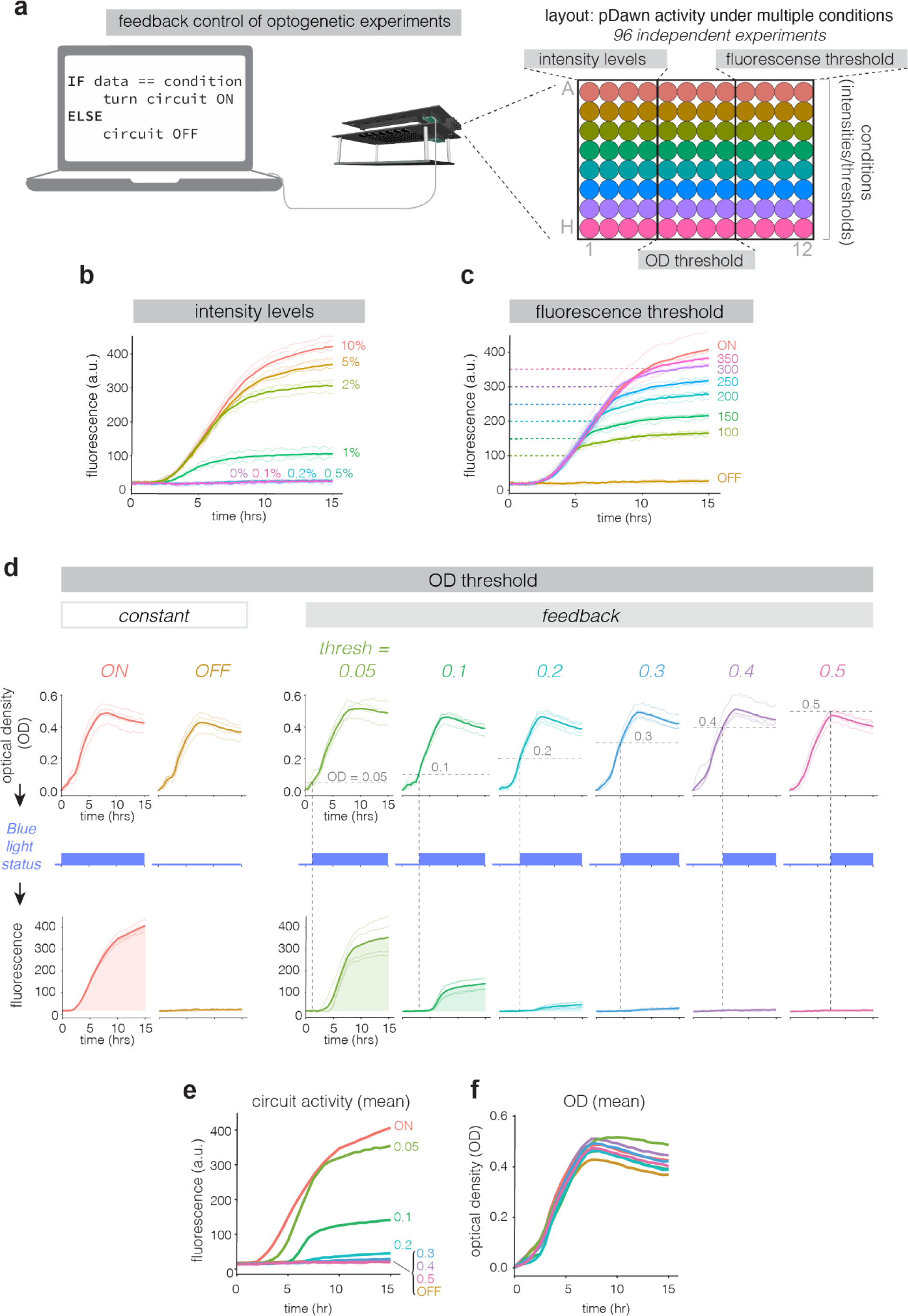
oPR allows computer-in-the-loop feedback control of bacterial behavior. **a** Diagram of “computer in the loop” feedback control of bacterial cultures, performed in 96 independently-controlled feedback experiments. The plate was divided into classes of experiments: constant light at different intensity levels, an fluorescence threshold at which blue light exposure is turned on, and an OD threshold at which blue light exposure is turned off. Each color of well represents a variation of a parameter within each feedback experimental class. **b** mAmetrine expression from bacteria stimulated with different light intensities. Each trace represents a single well at the corresponding blue light intensity. **c** mAmetrine expression in bacteria whose expression was halted once their fluorescence reached a predefined threshold. Dashed lines indicate the threshold for each trace. **d** OD (top row) and mAmetrine fluorescence (bottom row) in bacteria whose mAmetrine was optogenetically stimulated once their OD reached a pre-defined threshold. Dashed lines indicate the OD threshold. Mean fluorescence (**e**) and OD (**f**) are overlaid for clearer comparison of performance of each condition from (**d**). For (**b-d)**, each light trace represents a single well that was controlled based on its own state. bold traces represent the average of the four replicate wells per condition.

Despite the linear relationship, the magnitude of the oPR reading was approximately 1/2 of the Tecan reading (**Figure 5C**). Because Tecan readings are often used as a standard for OD readings, we derived a transformation for each well that adjusted oPR OD readings to reflect those of the Tecan (see **Methods** for full details). The equation for each well was then applied to oPR readings taken during a subsequent calibration experiment to generate “Adjusted oPR readings” which matched the amplitude of OD readings of the Tecan (**Figure 5E-G**). Importantly, oPR readings were consistent between separate experiments, meaning that one calibration experiment can be used to calculate adjustments that can then be applied to future oPR experiments. For a comparison between Tecan and oPR CVs at each OD, see **Supplementary Figure 7**.

### Using the oPR to stimulate and measure bacterial transcription and growth

To test the oPR’s ability to read bacterial growth and mAmetrine fluorescence, we generated *E. coli* that express mAmetrine under an arabinose-inducible promoter (**Figure 6A)**. We inoculated LB with these bacteria, plated in each well of a 96-well plate, and measured bacterial fluorescence and OD in the presence or absence of arabinose over 18 hrs in a 37 °C incubator. All live cell experiments were performed with 45 µL of parrafin oil deposited over 125 µL of culture samples. Paraffin oil prevented evaporation from the wells while permitting gas exchange and normal cell growth, consistent with previous work^22^ (**Supplementary Figure 8**). In all wells that received arabinose, the oPR detected mAmetrine fluorescence within the first two hours of arabinose addition, increasing and then plateauing throughout the experiment (**Figure 6B).** No fluorescence change was detected in the absence of arabinose. Optical density similarly increased and then plateaued in each well throughout the experiment (**Figure 6C**). In sum, these data demonstrate that the oPR can sensitively read fluorescence and absorbance over physiologically relevant regimes in bacterial culture, with low variability between wells. Moreover, these results show that the oPR is compatible with operation in bacterial incubators held at 37 °C and while shaking at 1000 RPM. Shaking dramatically enhanced growth of the bacterial cultures (**Supplementary Figure 9**).

Although the oPR optical configurations are optimized for mAmetrine, we also tested whether the device could be adapted to measure other fluorophores. By replacing the UV excitation LED with a blue LED (475 nm), the oPR could measure GFP expression in *E. coli* cultures (**Supplementary Figure 10)**. However, GFP, which was driven by the same promoter as mAmetrine in previous experiments, gave fluorescence signals that were ∼30% as strong as mAmetrine. This is likely because our emission filter more strongly attenuates GFP emission (507 nm) relative to mAmetrine emission (526 nm) (**Figure 1B**). Nevertheless, this experiment demonstrates that the oPR can be applied to detect fluorophores besides mAmetrine with appropriate changes to the LEDs and/or emission filters.

Next, we tested the oPR’s ability to optogenetically stimulate cells. We transformed cells with one of two plasmids — pDawn or pDusk^24^— that placed mAmetrine under blue light inducible transcriptional control. In pDawn, blue light stimulates reporter transcription that turns off in the dark, whereas in pDusk, blue light represses transcription that is otherwise constitutively active in the dark (**Figure 7A,B**). We seeded both strains in a single 96-well plate (48 wells for each strain). We then programmed the oPR to stimulate 24 wells of pDawn strain with blue light (pulsed with 3 seconds ON, 7 seconds OFF), while the remaining 24 wells each received no light. For the pDusk strain, the oPR was programmed to keep 24 wells in the dark while stimulating the remaining 24 wells in the same manner as for pDawn. Fluorescence and OD were measured every 30 minutes. The oPR successfully measured increasing mAmetrine fluorescence in all illuminated pDawn wells and dark pDusk wells over 10 hours of culture (**Figure 7C,D**, blue traces). Conversely, no fluorescence was detected in any dark pDawn well or illuminated pDusk well (grey traces), indicating that the oPR successfully confined optogenetic stimulation to only the desired wells. OD measurements successfully captured growth dynamics of both strains (**Figure 6C**). These data also demonstrate that UV light can be used to measure fluorescence without triggering a blue-light sensitive optogenetic tool, likely due to sparse illumination by the UV LED.

### All-optical feedback control in 96 independently-controlled experiments

The ability to simultaneously stimulate and measure cells allows for all-optical feedback control in 96 independent parallel experiments. To test this ability, we performed 96 experiments representing three classes of experiments that tested the main control knobs that could be implemented in custom feedback-controlled experiments (**Figure 8A**). The three classes of experiment were 1) constant open-loop blue light exposure with varying light intensities, 2) fluorescence-based feedback where optogenetic stimulation was halted once a particular mAmetrine fluorescence value was reached, and 3) OD-based feedback where optogenetic stimulation would only be applied after the culture reached a certain OD. For each class of experiment, we tested 8 distinct feedback parameters, and each parameter set was performed in biological quadruplicate, all on one microwell plate For experiments in class 1, we achieved graded expression mAmetrine between 0.5% and 10% of maximum blue light intensity (**Figure 8B**). For experiments in class 2, mAmetrine expression was successfully halted after fluorescence reached the designated level, with a small time lag due to inactivation kinetics of the transcriptional cassette, during which fluorescence continued to increase slightly before reaching a plateau (**Figure 8C**). For experiments in class 3, cultures triggered at higher ODs showed delayed mAmetrine expression and lower total levels of mAmetrine relative to cultures triggered at lower ODs (**Figure 8D-F**). These three experimental classes represent foundational operations that can be combined for more complex, custom feedback applications. Such custom functions can be written in the *FeedbackFuncs.py* file and can implement any mathematical process available in Python. Importantly, because of low well-to-well variability of stimulation and measurement in the calibrated oPR, 96 independent feedback-controlled experiments can be performed with low resultant variability between wells (**Figure 8B-D**).

## Discussion

The optoPlateReader (oPR) is a device for fully programmable optogenetic experiments, where stimulation, spectroscopic measurements, and feedback adjustments occur in an automated, pre-programmed manner, in 96-well plates. Because there are no moving parts and no wires other than power cables and USB cables, the oPR is robust to mechanical perturbations, allowing it to be used on shakers and within incubators. We showed that the oPR can read fluorescence and OD from bacteria with high sensitivity and low variability between the 96 well positions. The high precision measurements allowed direct comparison of different wells in the same plate, and further enabled computer-in-the-loop feedback control, where the optogenetic stimulus of each well can be dynamically updated based on its current state. We also developed software that integrates and controls all components of this device, including a graphical user interface that allows easy programming of complex optical reading, writing, and feedback parameters for each individual well. Importantly, the oPR is an open source device that can be built with no prior expertise in electronics within ∼6 hours, at low cost, and can be readily modified for custom applications.

We overcame two main obstacles in our hardware design. The first was in optimizing the optical components and their orientation to allow sensitive OD600 and fluorescence measurements. In particular, the compact placement of the photodiode and the adjacent UV LED could lead to contamination of the fluorescent signal with the excitation light. We overcame this challenge in three ways. First, we used a photodiode that was mounted on the opposing side of the PCB and faced the sample through an aperture. Second, we selected a photodiode with a low responsivity in the UV range, providing a measure of filtration of the UV LED light. Finally, we included a plastic emission filter film that further blocked light in the UV range but passed light between 470 nm and 650 nm, permitting OD600 measurements and mAmetrine emission detection. These optimizations were enabled by selection of the long-Stokes-shift fluorescent protein mAmetrine, which provides sufficient separation in the excitation and emission spectra to allow effective filtration of excitation light with inexpensive filters while minimally affecting fluorescence or OD detection. A second important design challenge was in the variability among the three LEDs and one photodiode. After device assembly, we measured up to ∼20% variability between the 96 components of each type. Such variability could further compound, for example if UV LEDs with high variance are used to stimulate fluorescence that is measured by photodiodes with high variance. Such measurements would have high uncertainty, challenging interpretation and making comparisons of values from different wells impossible. We overcame this limitation by 1) designing 3D adapters for optimal and consistent alignment between oPR components, and 2) developing careful calibration protocols, first for the photodiodes, and then using the calibrated photodiodes to calibrate remaining LEDs. These measures were essential to obtaining the tight correspondence of fluorescence values between wells of the same conditions, and for the robust OD measurements that enabled precise OD-based feedback control. If further reduction of technical variability is required, readings from replicate wells can be averaged together during the experiment via the “grouping function” found in the GUI (See **oPR Repository Usage Manual**).

The ability to perform 96 parallel feedback experiments represents the biggest advance enabled by the oPR. Such control could be highly advantageous for fields like metabolic engineering, where optogenetic transcriptional control can dynamically optimize flux through engineered metabolic pathways^25^. Although the parameter space for such optimizations is vast (spanning different intensities, waveforms, duty cycles, durations, etc), the ability to automate ∼100 such experiments simultaneously and without user intervention during the experiment allows dramatically faster sampling of these parameters. Furthermore, feedback control algorithms could allow the oPR to optimize stimulation protocols on the fly, outputting the stimulation parameters it used to achieve the predefined target state. Moreover, because of the small footprint and low cost of the oPR, multiple devices could be operated in parallel, further increasing throughput.

Although we characterized the oPR for measuring growth, fluorescence, and for optogenetic stimulation of bacterial cultures, the device and its submodules could be applied more broadly. Similar experiments could be performed for example in yeast, mammalian cells, or even cell-free systems, though the electrical circuitry that sets the sensitivity and dynamic range would likely have to be optimized for each application separately. In addition, the optoReader module could be used as a standalone device, similar to an existing 96-well phototransistor array that measured absorbance^19^, but with the added capability of fluorescence measurements. Alternatively, a well-calibrated optoReader could be used to rapidly calibrate the optoPlate-96 or other LED arrays used for optogenetics, which have a typical variation of 10-20% in intensity between LEDs after assembly^2,27,28^ and thus require calibration for precise greyscale optogenetic control.

Several future improvements could further empower experiments with the oPR. Optimization for fluorescent proteins other than mAmetrine, or even to allow reading of multiple fluorescent proteins, would expand experimental capabilities. Such modifications should be feasible with high-quality emission filters like those found on fluorescence microscopes, though at increased cost. The oPR also has exposed circuitry which has the potential to become corroded, for example as a result of accidental spills of culture media. Thus a physical barrier to protect the circuitry would increase its robustness, though we note that no spillage or loss of functionality was observed despite shaking during all live-cell experiments. In addition, the oPR circuitry could be enhanced to allow wireless communication with the computer, and also to increase sensitivity and dynamic range of light detection.

In sum, the oPR is device for high-throughput, feedback-enabled optical reading and writing in cells, and it achieves such sophisticated experiments in a compact, inexpensive, and open-source manner, opening new horizons for optogenetic and cybernetic interfaces with biological systems.

## Acknowledgements

We thank the George H. Stephenson Foundation Educational Laboratory & Bio-MakerSpace for providing space and resources necessary for completion of this work. This work was supported by the Bradley Gabel Memorial Fund (to S.D., L.S., J.H., G.C., and G.L.), the National Institutes of Health (R35GM138211 for L.J.B and D.G.M., R01NS101106 to B.Y.C.), the National Science Foundation (CAREER CBET 2145699 to L.J.B., CAREER MCB 1652003 to B.Y.C., CAREER CBET 1751840 to J.L.A., GRFP to W.B. and G.H), the U.S. Department of Energy (Office of Science, Office of Biological and Environmental Research, Genomic Science Program under award number DE-SC0019363 and DE-SC0022155 to J.L.A) and Singapore’s STAR Fellowship to S.M.

## Author Contributions

W.B., D.G.M., J.L.A, B.Y.C.,L.J.B conceived study. S.D., W.B., D.G.M, G.L., J.H., G.Q., G.L., L.S., G.H., S.M., M.S.M., M.P., S.M., J.L.A., B.Y.C., L.J.B. designed hardware structure and component layout. S.D., W.B., D.G.M, G.L., J.H., G.Q., G.L., L.S., G.H., S.M., L.J.B. constructed hardware, performed experiments, and analyzed data. S.D., G.L., J.H., W.B., D.G.M, developed software. S.G.M., J.L.A., B.Y.C., L.J.B. supervised and advised work. S.D., W.B., L.J.B. drafted manuscript and figures. All authors contributed to editing.

## Supplementary Figures

**Supplementary Figure 1.**
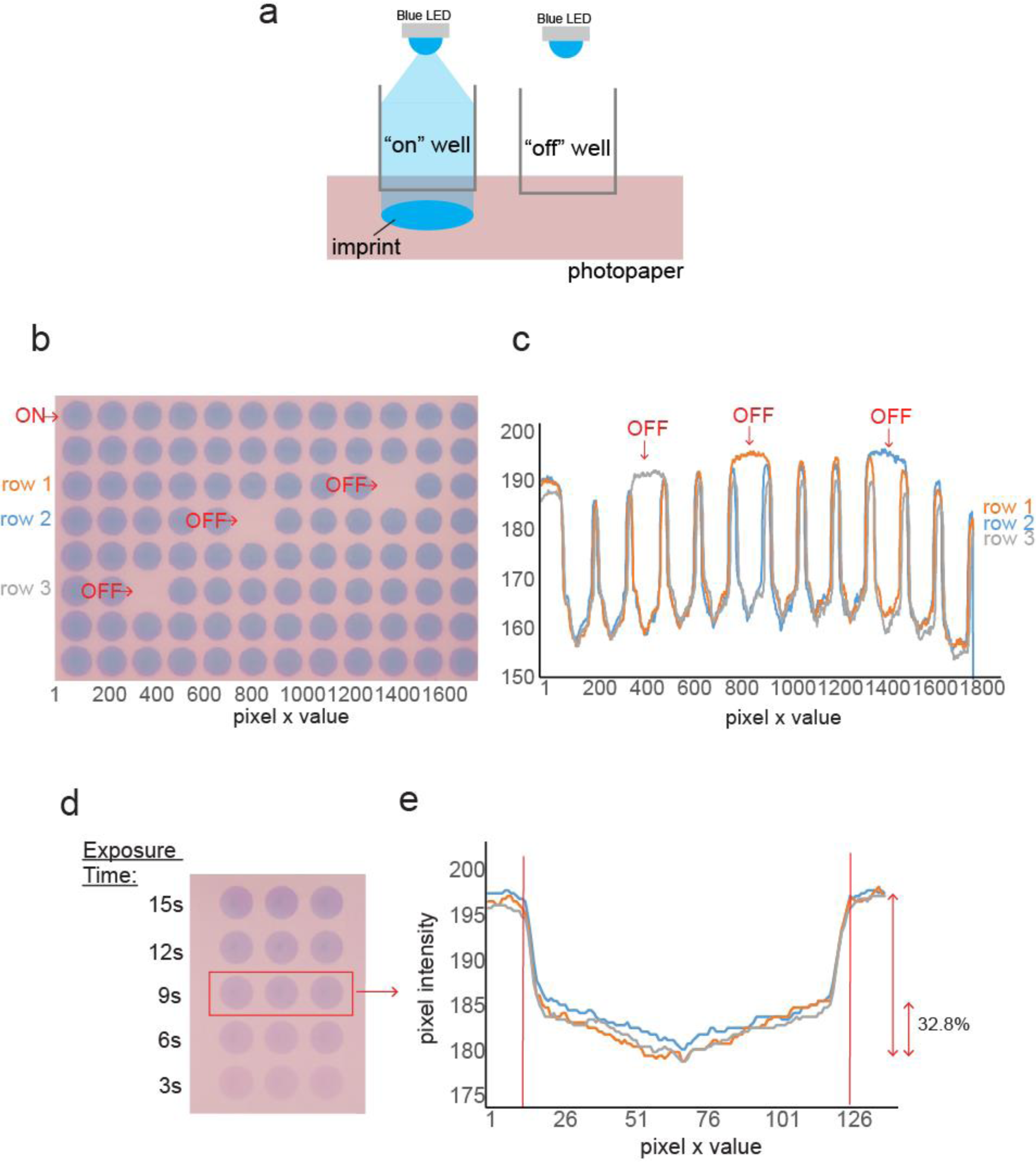
Insulation of light between wells and homogeneity within wells. **a** Photopaper allows for imprinting and recording of light exposure underneath a 96 well plate from the blue LEDs^9^. **b** All wells of the optoPlate’s blue LEDs were set to maximum except the 3 indicated as “OFF.” The optoPlate was mounted on a 96 well plate with photopaper placed below the plate. No imprint was observed in OFF wells, despite all surrounding wells being set to maximum intensity for 5 minutes. This indicates that light spread between wells is negligible. **c** Quantification of the intensity across each row of wells of the exposed photopaper in **(b)** shows that OFF wells do not display detectable imprints from light exposure despite neighboring wells being set to maximum intensity. Data represents a line profile of the pixel intensity across the center of each row of wells. **d** Photopaper can also be used to analyze light uniformity within wells. Triplicate wells were exposed to blue light for varying times in order to find a non-saturating exposure time. 9s was chosen for quantification. **e** Line profile of the pixel intensities across the center of each well from the 9s exposure condition from **(d)**. Blue light intensity varies by 32.8% from the center to the edge of the wells (indicated with red lines). However, this non-uniformity is can be overcome either by 1) stimulating with strong blue light such that even the weaker light at the periphery is saturating for the optogenetic protein, or 2) shaking the cultures during growth, homogenizing the effective light intensity that the bacteria experience over time.

**Supplementary Figure 2.**
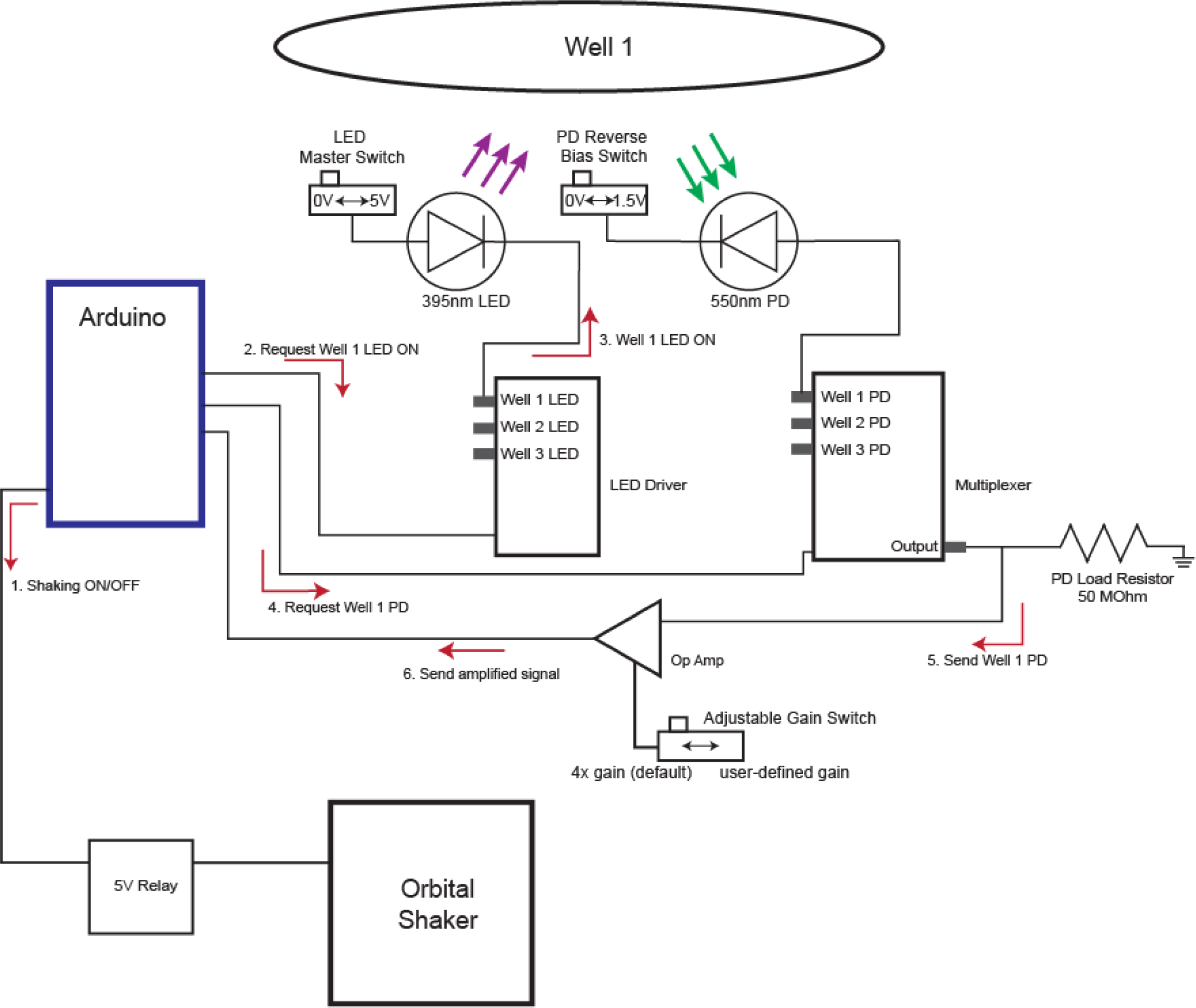
oPR circuit diagram for fluorescence readings in a well. To acquire fluorescence readings for a well, the following sequence of events occurs: 1) Shaking is paused during measurements. 2) The oPR Arduino sends a signal to the LED drivers requesting that the UV LED for that well be switched on; 3) The LED driver switches on the LED for that well while leaving all other LEDs off. The LED 5V switch must be set to the 5V position for LEDs to turn on. This switch ensures that the LEDS are only operational when desired to prevent inadvertent illumination on samples or investigators. 4) The Arduino then sends a command to the multiplexer to select the photodiode (PD) for that well. The PD signal can be altered by changing the Reverse Bias switch. 0V (used in this study) allows detection of lower intensity signals by reducing the background signal produced by the photodiodes. 1.5V allows for detection of higher intensity signals by raising the point of signal saturation. This is because a greater maximum voltage can be produced by the photodiode than is possible without a reverse bias. However, this also causes a higher background signal when no fluorescence is present. 5) The signal from the PD is then sent to an amplifier that multiplies the signal by an adjustable gain. The Adjustable Gain switch can be set to 4x (used in this study) or a user defined gain (by adding a resistor of a desired value as labeled in the component diagram). 6) The final signal is sent to the Arduino for processing. A similar sequence of events takes place for OD readings, with additional communication with the optoPlate to coordinate illumination of the OD LED.

**Supplementary Figure 3.**
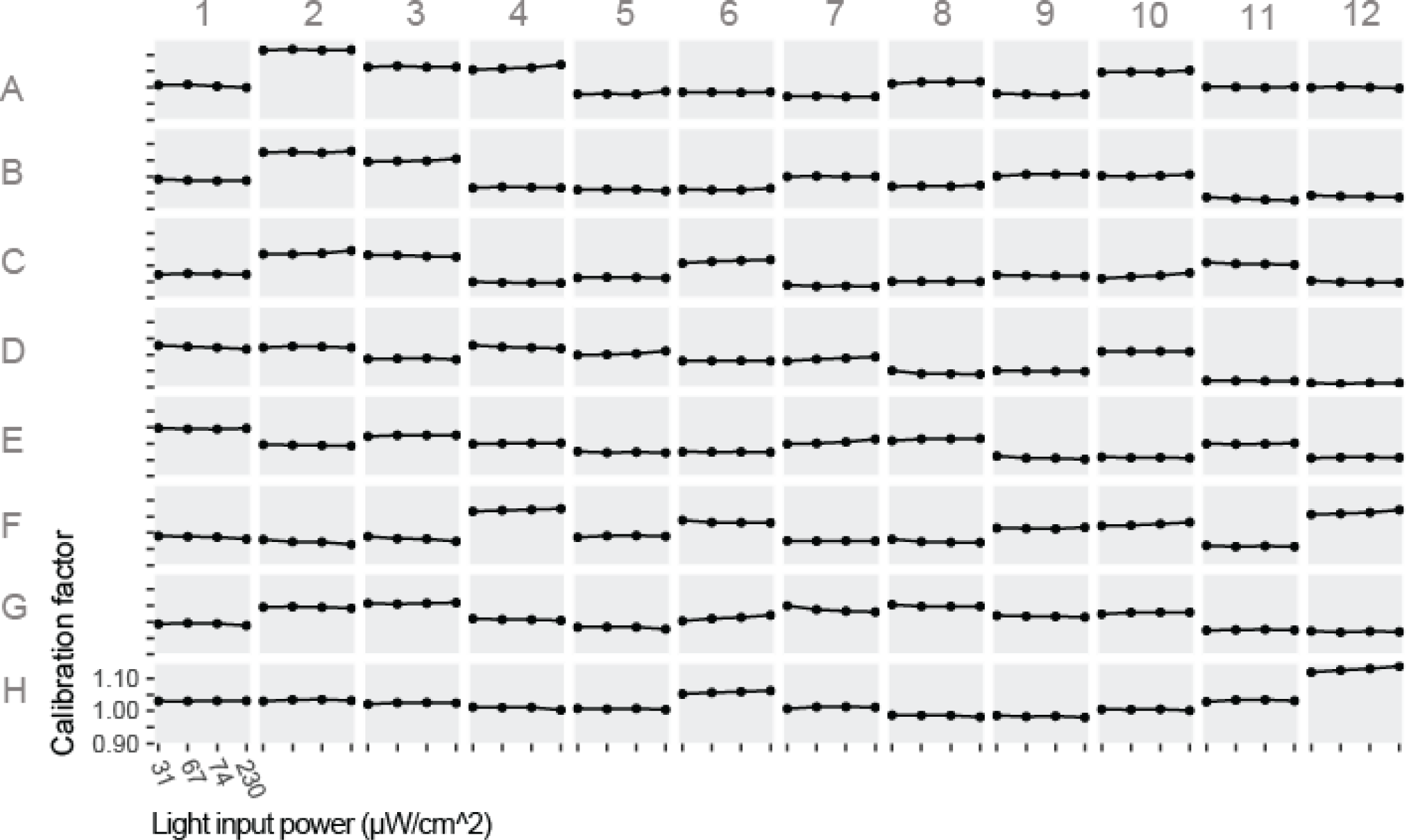
Photodiode calibration factors are constant at different light input intensities. The calibration factor for each input intensity was calculated separately for each well at 4 different light intensities. In general, calibration factors were independent of input light power.

**Supplementary Figure 4.**
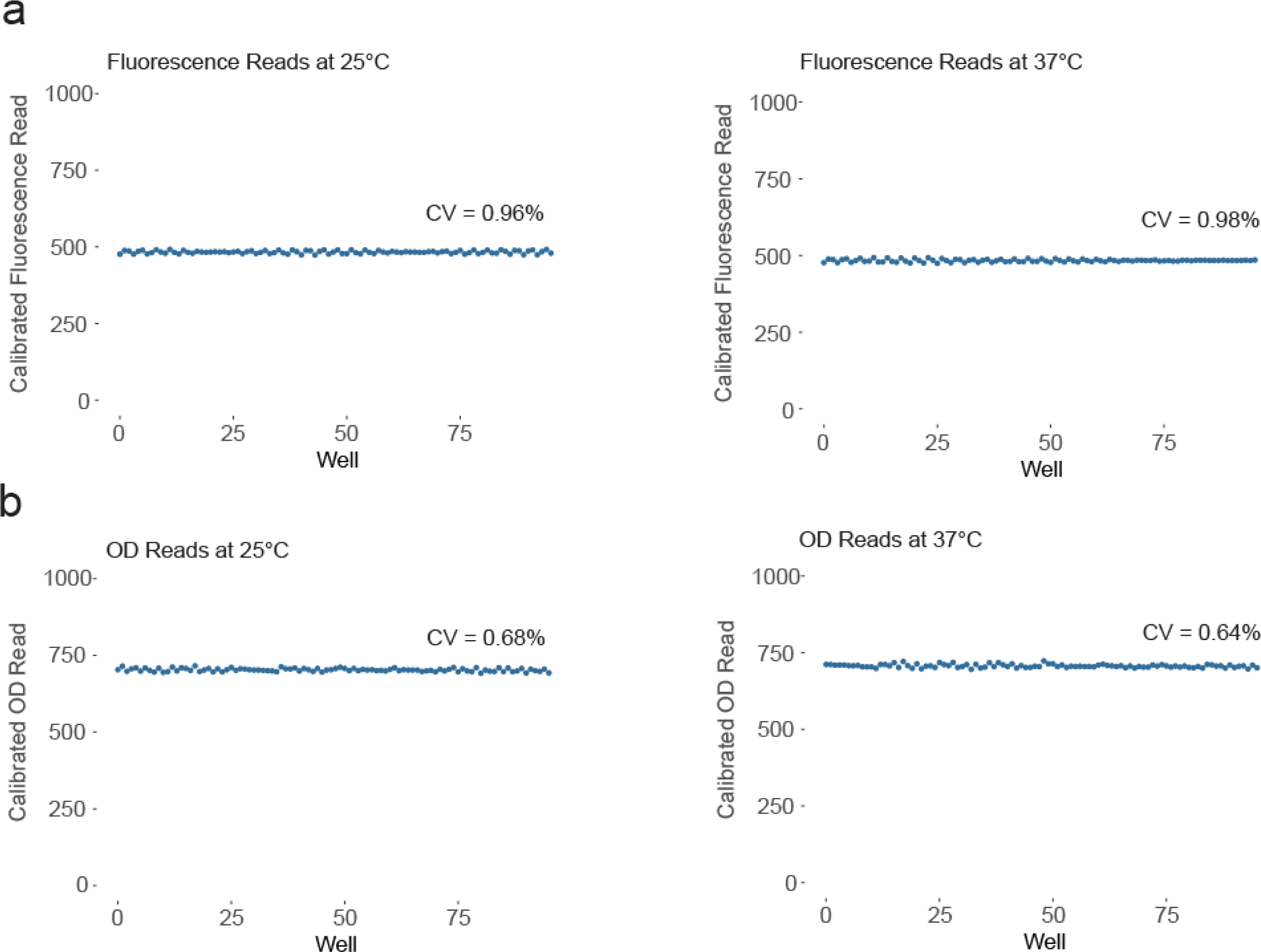
Calibrated measurements do not vary as a function of temperature. **a** Calibrated fluorescence measurements were taken at 25°C using 40 μg/mL of lucifer yellow per well inside of a cell culture incubator. The cell culture incubator was then allowed to equilibrate at 37°C and measurements were repeated using the same calibration values. There was no significant change in measurement amplitude or variability of measurements between the two temperatures. **b** The same process was repeated for OD LED intensity with wells containing LB and again no significant temperature-dependent differences were observed.

**Supplementary Figure 5.**
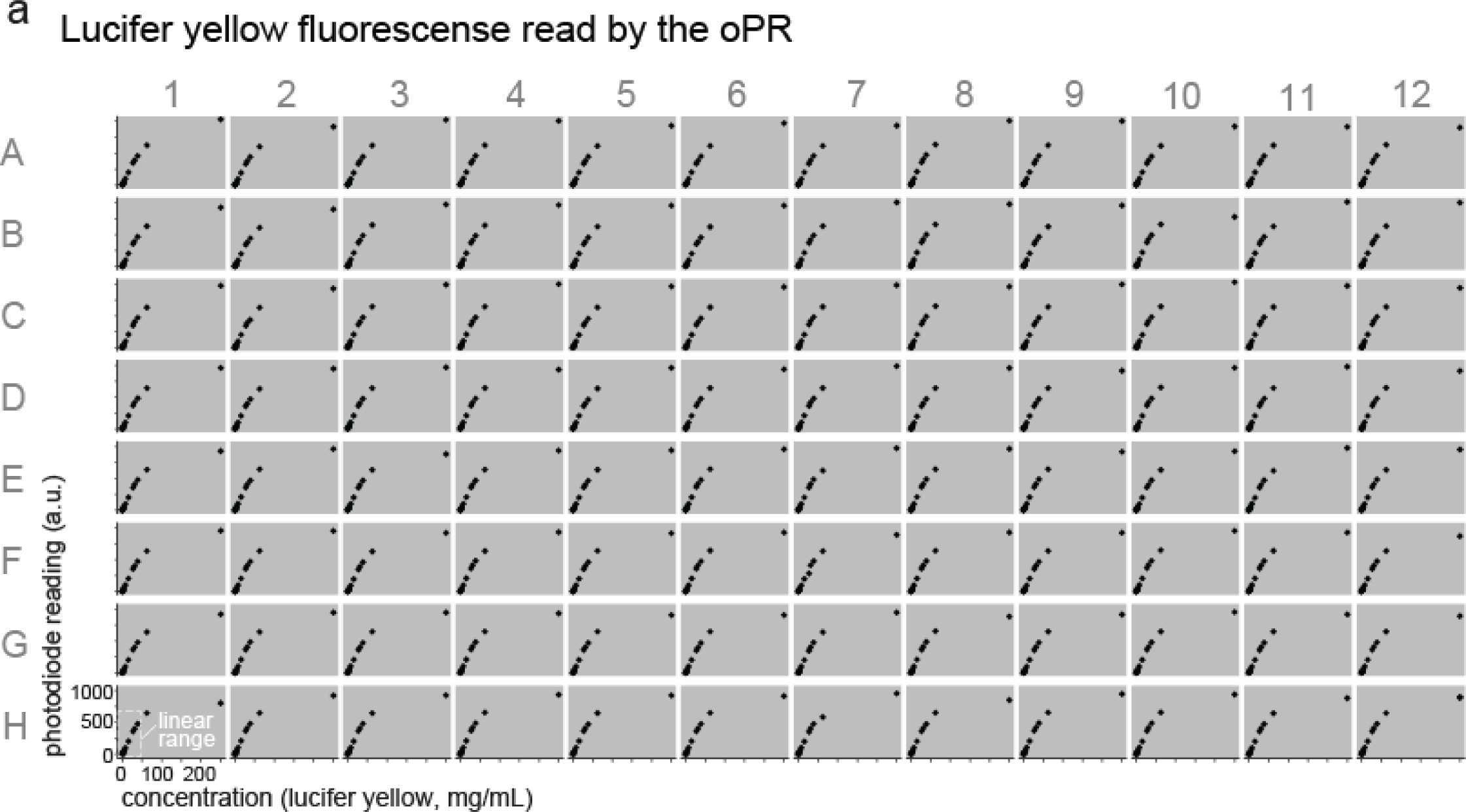
Full range of photodiode fluorescence characterization. Fluorescence measurements of lucifer yellow dye were taken over a range of concentrations (2-250 µg/mL) in each of 96 wells in a calibrated oPR. Concentrations from 2-40 µg/mL are reproduced from Figure 4.

**Supplementary Figure 6.**
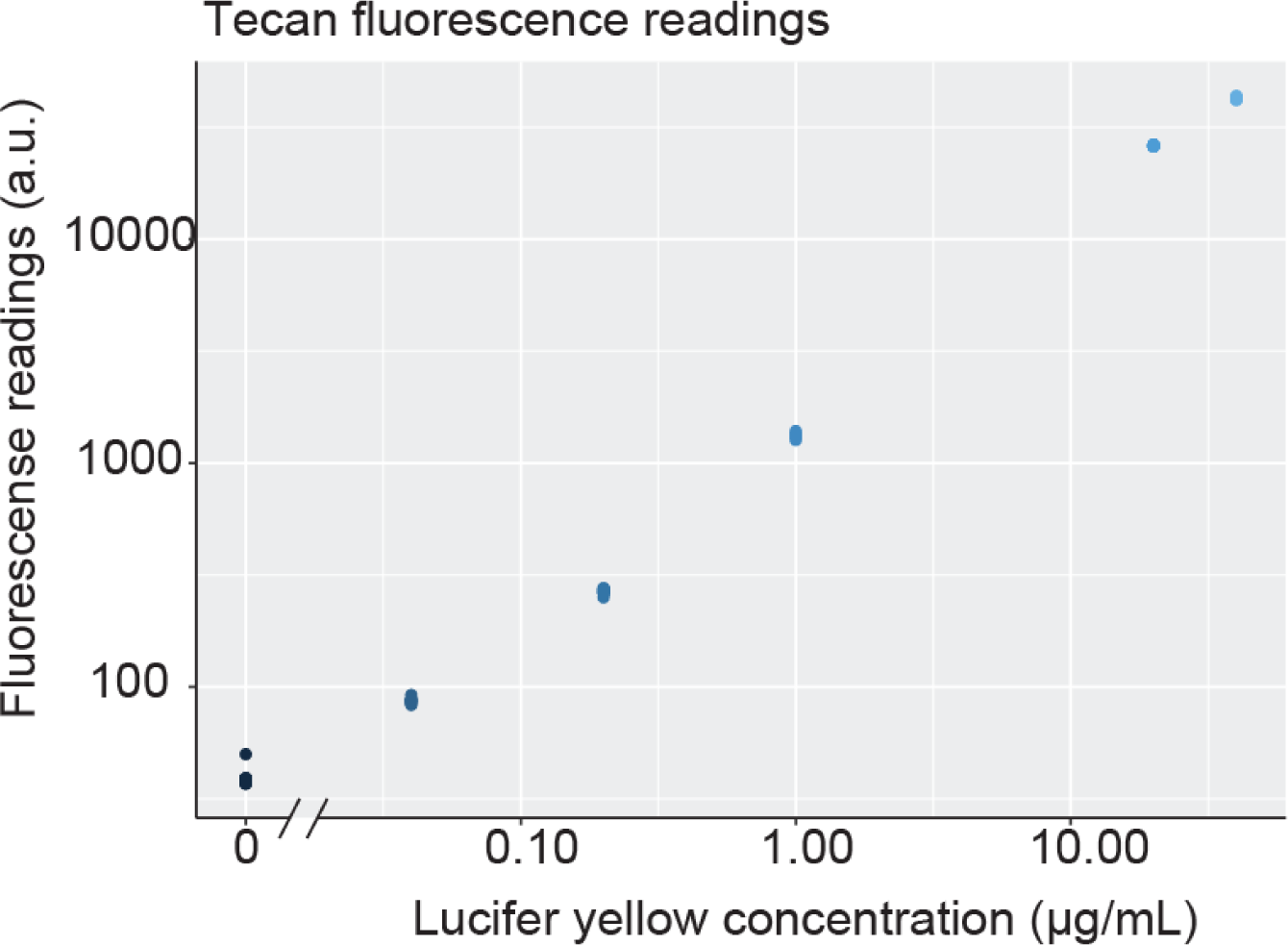
Tecan limit of detection experiments. Lucifer yellow was diluted to several concentrations and measured using the Tecan Infinite M200. Based on this data, the limit of detection was calculated to be ∼10 ng/mL. Each point represents the read from 1 well of lucifer yellow at the specified concentration.

**Supplementary Figure 7.**
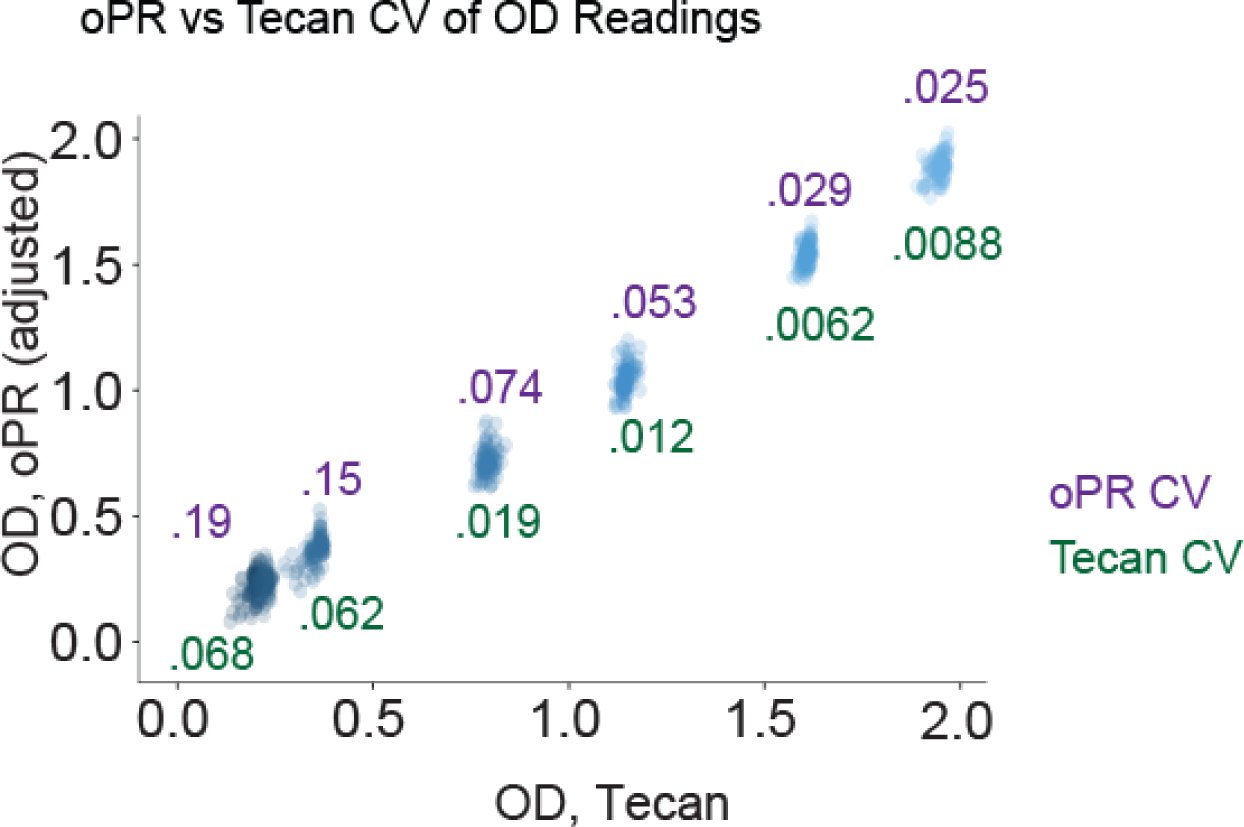
Coefficient of variation for oPR vs Tecan OD readings. CV values were calculated for data shown in **Figure B**. CVs displayed above data points correspond to oPR measurements (magenta) at the corresponding concentration. CVs displayed below data points correspond to Tecan measurements (green).

**Supplementary Figure 8.**
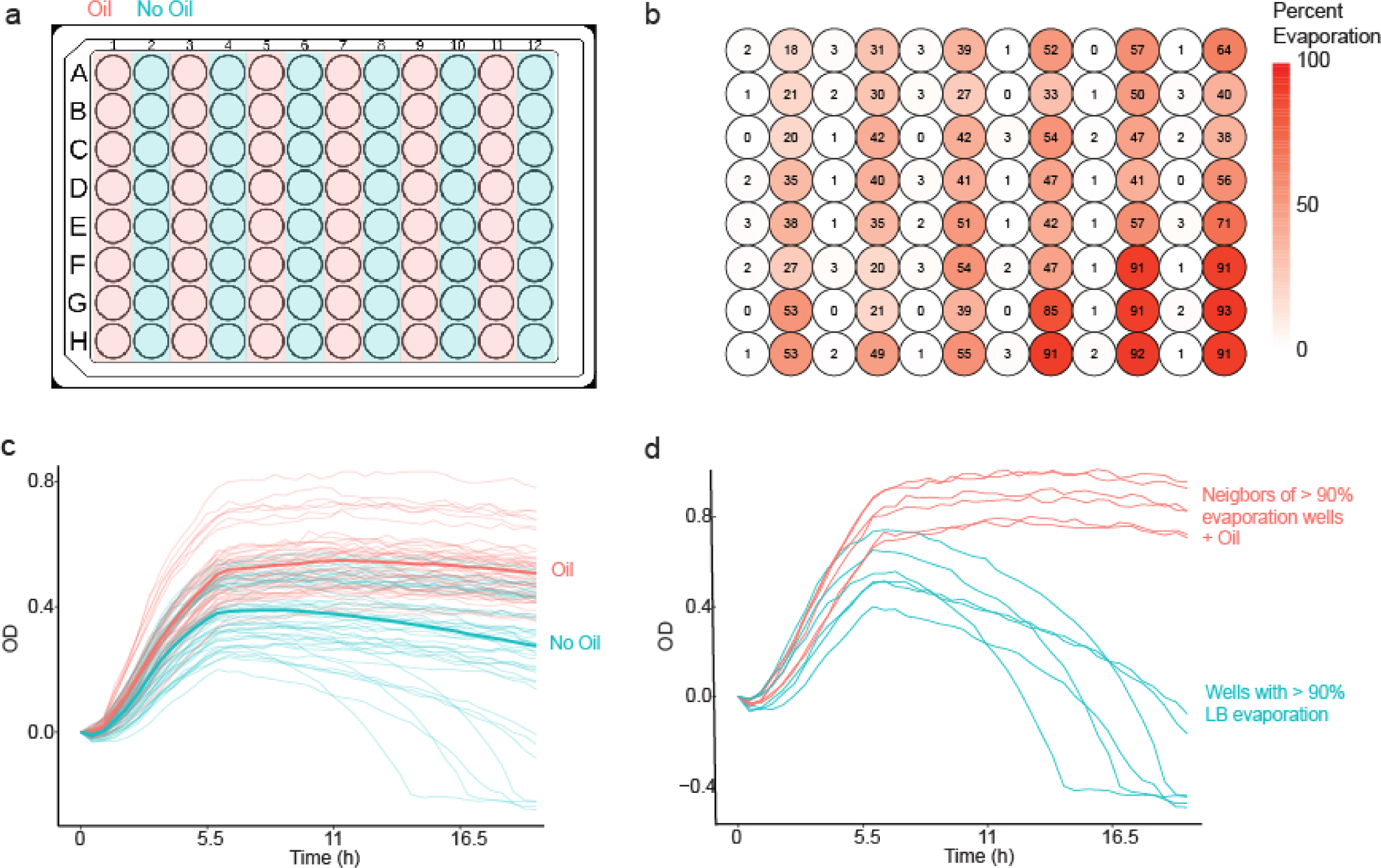
Paraffin Oil prevents evaporation. **a** Bacteria were cultured in a 96-well plate with and without a 45 µL layer of paraffin oil for 18 hours of culture and measurement on an oPR with shaking at 1000rpm. **b** Percent LB evaporation from each well after the experiment described in (**a**). **c** OD readings over time from the experiment described in (**a**). Wells without paraffin oil exhibit an initial increase but subsequent decrease in OD readings, likely due to the fact that evaporation caused settling of dry residue on the outer edges of the well, creating an optically clear path through the center of the well, allowing greater light transmission than the initial liquid LB. By contrast, wells with paraffin oil exhibit a monotonic increase and plateau at ∼ 6 hrs. **d** Examination of wells with most extreme evaporation compared to their neighbors with paraffin oil. Wells with >90% evaporation show a major decrease in OD readings over time (blue) while their direct oil-coated neighbors showed normal growth (red).

**Supplementary Figure 9.**
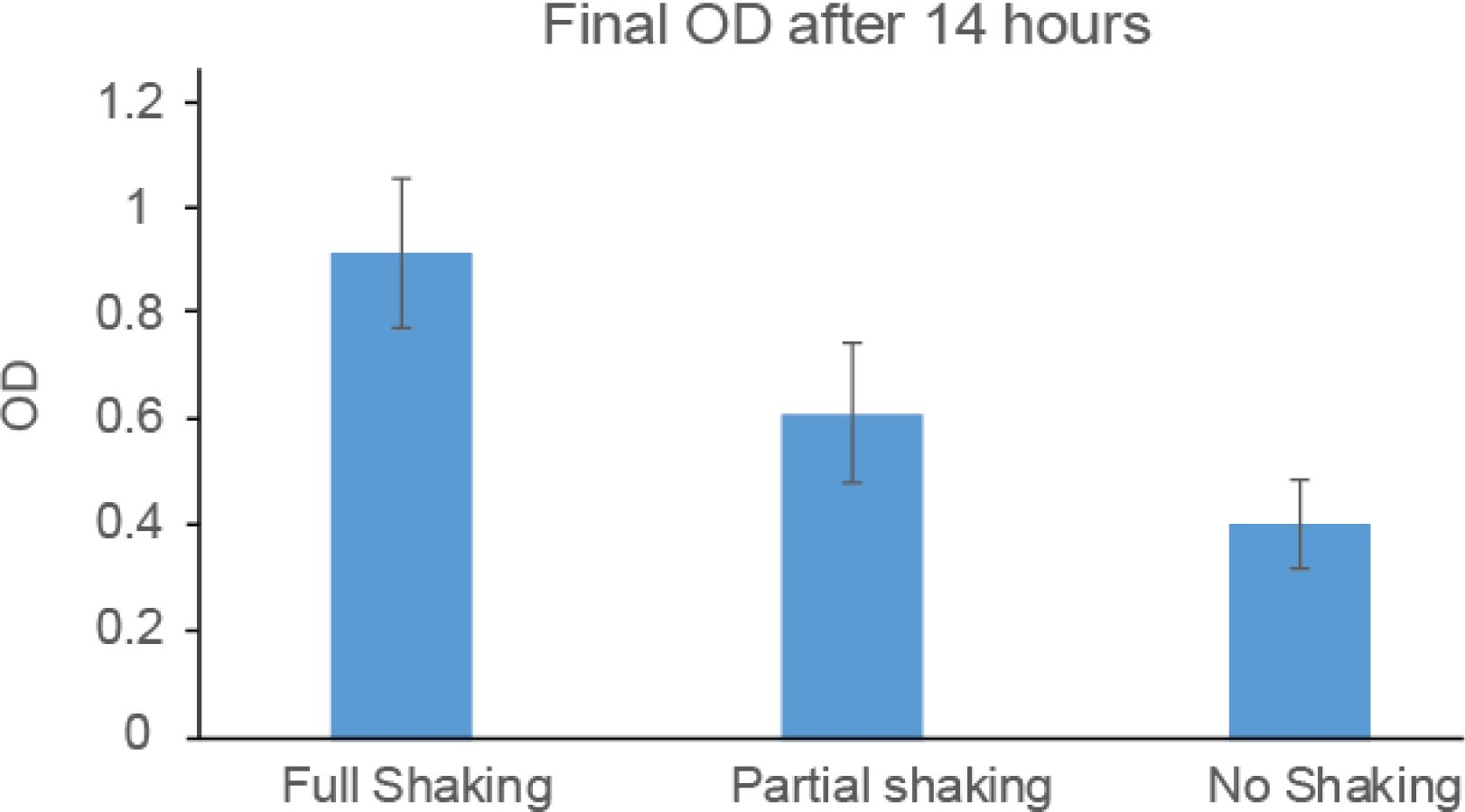
Shaking effect on culture OD. Bacterial cultures were grown in 96 well plates for 14 hours with either constant shaking, 2 minutes of shaking every half hour, or no shaking. The final OD was measured, showing that increased shaking frequency led to higher final ODs. Each bar represents the mean of 96 wells with error bars representing 1 s.d.

**Supplementary Figure 10.**
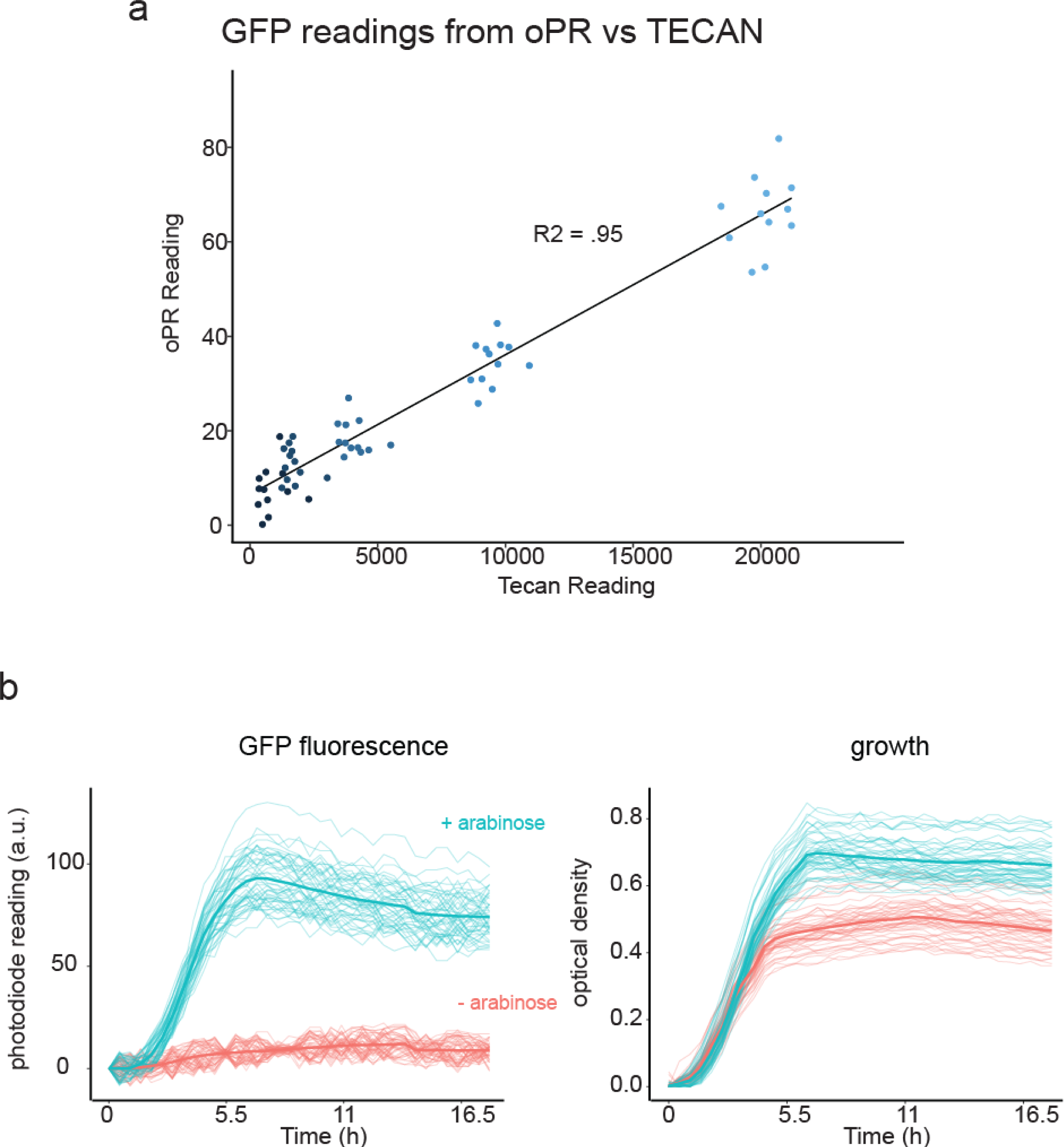
oPR detection of GFP in live cultures. **a** Bacteria transformed with an arabinose inducible GFP expression plasmid were grown to saturation in 2 mL of LB + arabinose and diluted in a series of 2x dilutions. Each dilution was seeded in one row of a 96 well plate, and GFP intensities were read with both an oPR and Tecan. The oPR was able to detect GFP using a 475 nm excitation LED, and increases in fluorescence were linear with increases in fluorescence as measured by a Tecan plate reader. **b** Arabinose inducible GFP cultures were grown with or without arabinose for 18 hours, demonstrating the ability of the oPR to detect GFP expression during live time course experiments..

## Methods

### oPR Repository

All files including 3D printed adapters, PCB designs, Arduino and Python scripts, assembly manuals, and usage manuals can be found in the attached **oPR repository**.

### Circuit Design and CAD

All circuits and PCBs were designed using Kicad version 5.1.6-1. All CAD files for 3D printed adapters were designed using the Rhino software. All files are freely available in our online oPR repository.

### oPR construction

oPR construction was achieved using standard PCB assembly protocols. The optoPlate was assembled as previously described^1^ but with bi-color LEDs (Wurth Elektronik 150141RB73100) and a 589 nm OD LED (Broadcom HSMA-C380) used as the two LEDs for each well. The optoReader is a 2-sided PCB and was assembled as follows. Briefly, solder paste (Chipquik SMD4300AX250T3 183 °C) was deposited through a mask onto the top side of the PCB, which was fabricated by JLCPCB. Components were then placed and the PCB was baked in a toaster oven at 190 °C until solder paste became visibly melted. The PCB was allowed to cool to room temperature. Then, solder paste with a lower melting point (Chipquik TS391LT50 138 °C) was deposited via a mask on the bottom side of the PCB, components were placed, and the PCB was baked in a toaster oven at 150 °C until solder paste became visible melted. Detailed component placement diagrams can be found in our online repository (See first section of **Methods**). 2 layers of Rosco #312 filter sheets were used as emission filters for fluorescence detection. Each filter layer was cut using a Silhouette Cameo craft cutter using (cutting files are found in the oPR repository). The device acrylic base plate was laser cut using the laser cutting file also found in the file repository.

### Experimental Conditions

For fluorescence calibration, a stock of Lucifer Yellow dye (Invitrogen, L453) was diluted in DI water to a concentration of 1 mg/mL, and was further diluted in water to the concentrations indicated. oPR fluorescence measurements were taken with 200 µL of dye solution in each well. For OD testing, 1.1 µm plastic beads (Millipore Sigma #LB11-1ML) were used to create standard dilutions for OD calibration. For each bead dilution, dilutions were added to a 96 well plate and moved directly to the oPR where OD was measured. The plate was then transferred to the TECAN, shaken for 10 seconds at 432 rpm, and OD was measured. For oPR experiments using bacterial samples, LB medium with the appropriate antibiotic was inoculated with bacteria, and 125 µL of the inoculated medium was seeded in each well of a 96-well plate and coated with 45uL of paraffin oil. 100 µL DI water was added to the space between wells to suppress sample evaporation. OD LEDs were calibrated at the start of each experiment with the starting (T = 0) cultures, representing the blank (max transmission). The blue LED intensity was programmed to stimulate the sample at an intensity of 1000 (∼25% of max intensity) with a 30% duty cycle, stimulated for 3 second pulses with 7 second intervals of darkness between pulses. Pulsing was used to minimize potential phototoxicity. All experiments were carried out in a 37 °C incubator with shaking at 1000rpm.

### Generation of pDawn/pDusk-mAmetrine E. coli

pDawn and pDusk plasmid with a GFP reporter were a gift from Andreas Moeglich (Addgene plasmids #43796 and 43796). Arabinose inducible mAmetrine (pBad-mAmetrine) was a gift from Robert Campbell (Addgene plasmid # 18083). mAmetrine was inserted in place of GFP in both pDawn and pDusk plasmids via PCR and Gibson assembly (NEB HiFi mix). Using the same method, mAmetrine was replaced with GFP in the pBad-mAmetrine plasmid to create arabinose inducible mAmetrine. Constructs were transformed into NEB DH5a competent *E. coli*.

### Software

To operate the optoPlateReader, a Python program communicates with two Arduinos, one on the optoPlate and one on the optoReader. All software files and usage instructions can be found in our online oPR repository. Prior to the start of each experiment, the *optoReader.ino* and *optoPlate.ino* Arduino sketches are uploaded individually to the respective Arduinos via USB. The *optoReader.ino* Arduino sketch operates the optoReader Arduino and coordinates photodiode measurements and UV/excitation control. The *optoPlate.ino* Arduino sketch operates the optoPlate and coordinates the blue and red (OD) LEDs. Next, the *Protocol.py* and *GUI.py* scripts are opened on a central computer. The Arduino ports through which serial communication occurs are hardcoded in the *Protocol.py* script and are edited accordingly where specified. The *Protocol.py* program communicates with the two Arduinos. This program translates the user defined LED illumination intensities, timing, reading intervals, and experimental duration into commands for the Arduinos. It also receives photodiode measurements from each well, and it calculates user-defined feedback functions and updates the oPR protocols in real time. The final step before running the experiment is to run *GUI.py*. This script opens a graphical user interface that provides a user-friendly method to define the experimental parameters that *Protocol.py* then feeds to the Arduinos. In the GUI, the user defines the groups of wells that receive the same experimental conditions and then defines those conditions, including the stimulation light intensity, duty cycle, and feedback functions. The user then specifies measurement parameters for fluorescence and OD readings, including the frequency of measurement and the number of measurements to average for one reading. After defining these conditions, the experiment is started by pressing the “Run Plate” button. The two Arduinos remained connected to the computer through USB cables for the duration of the experiment. For more details on software, see the usage manual found in our oPR repository.

### Calibration

Calibration of the optoPlate components was essential to obtain precise measurements between all 96 wells by ensuring consistent stimulation intensities and photodiode measurements. We first calibrated the photodiodes to ensure consistent light measurements, and then we used the calibrated photodiodes to calibrate all LED components. While photodiode calibration must be performed in the manner we describe below, LED calibration can be performed through the GUI, which automates the steps described below.

To calibrate the photodiodes, a photo light box (Havox Hpb-40d) was used as a uniform light source to ensure that each photodiode was illuminated with the same intensity of light. An external light meter (Thorlabs, PM100D) was used to ensure uniformity across the illumination region (< 3% variation). The optoReader was placed in the center of the light box, connected to the computer, and plugged into the power supply. No adaptor or emission filter was used. The *optoReader_Manual.ino* script was uploaded and run to collect 100 readings per well for 96 wells. The following procedure was used to determine the calibration factor. First, the mean of the 100 readings was calculated for each well. Then, the calibration factors for each well (c_i,pd_) were calculated by dividing the minimum reading of all wells by the average reading for that well:

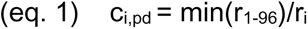

Four calibration factors were calculated in this manner using four different illumination intensities (31,67, 74, 230 µW/cm^2^) to ensure that the photodiode calibration was consistent across light intensities. Differing illumination intensities were achieved by placing diffuser film between the light source and the oPR. The final calibration factor for each photodiode was calculated by averaging the four calibration factors per well (calculated for each irradiance value). However, calibration factors did not vary as a function of illumination intensity (**Supplementary Figure 3**). The photodiode calibration factors were manually input to the Python script, *Protocol.py*, during initial set-up of the oPR.

LED intensity on the optoPlate and optoReader is controlled by the TLC5947 LED driver chip (Texas Instruments). Each chip controls intensity of up to 24 LEDs through 4095 intensity levels using pulse-wave modulation (PWM). oPR LEDs were calibrated by calculating correction factors that were then applied to adjust the PWM settings for each LED.

The stimulation (blue) LEDs were calibrated in the fully assembled oPR but without the excitation filter film, which would have attenuated the blue light. An empty black, clear bottom 96-well plate (Greiner, #07000166) was placed between the optoReader and optoPlate in order to calibrate the LEDs through air. To calibrate, the script *optoReader_Manual.ino* was uploaded to the optoReader, and the script *optoPlate_Manual.ino* was uploaded to the optoPlate. These scripts set the blue LED intensity to a value of 2000 to begin the calibration. The optoReader reported the average of 100 readings per well of the 96 well array. The coefficient of variation (CV, standard deviation/mean x 100%) of the readings was calculated. From these initial readings, a calibration factor for each well (c_i,bl_) was determined with the following formula:

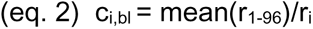

where r is the mean reading from the optoReader. To test the calibration, the blue light intensities for each well were multiplied by their calibration factor. These values were updated in the *optoPlate_Manual.ino* script, which was uploaded to the optoPlate. A new set of readings was obtained, and the CV was calculated. This process was iterated until the CV reached < 1%, or up to 3 rounds (no decrease in CV was observed after 3 rounds). The final blue LED calibration factors were saved to a .csv file through the GUI and manually added to the *optoPlate.ino* Arduino script at the initial set-up of the oPR.

The optical density (OD, red) LEDs were calibrated in a similar manner to the blue LEDs above, with a few key differences. First, the OD LEDs were calibrated such that the OD readings were consistent between wells. This means that calibration normalizes not just LED intensity, but also slight variations in alignment and light scatter that can occur between wells. To achieve this, calibration must be performed with liquid in each well, since the optical properties of the sample liquid can differentially scatter the red light. Second, OD LED calibration was performed at the beginning of every experiment to account for differences in liquid volume, clarity, meniscus, etc. from experiment to experiment (other LEDs were only calibrated once after initial construction of the oPR device). To begin calibration, a black, clear bottom 96-well was filled with 200 µL of LB in each well, and the full oPR was assembled. The script *optoPlate_Manual.ino* set the OD LEDs to an initial brightness setting of 500. The optoReader reported the average of 100 readings per well. The CV of the measurements was calculated, and the calibration factors for each well (c_i,OD_) were calculated using the following equation:

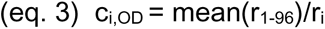

As before, the calibration factors were used to adjust the OD LED intensity settings. These adjusted values were then uploaded to the optoPlate, and new measurements were taken. This measurement/adjustment cycle was performed iteratively until the CV of the readings reached < 1%, or for up to three rounds. The final OD LED calibration factors were saved through the GUI as a .csv file and were referenced by the *Protocol.py* program throughout the experiment.

The UV LEDs were calibrated based on the measured oPR fluorescence of a uniform concentration of fluorescent dye, lucifer yellow, which has excitation and emission spectra comparable to mAmetrine. To begin calibration, 200 µL of 1000 µg/mL lucifer yellow diluted in water were deposited in each well of a clear-bottom, black-walled plate, and the oPR was fully assembled. The script *optoReader_Manual.ino* was modified to set the UV LEDs to a maximum intensity setting (4095), and the script was uploaded to the optoReader Arduino. The optoPlate was mounted over the well-plate as usual, but no script was uploaded to the optoPlate Arduino since none of its components were required for UV LED calibration. Photodiode measurements were recorded as before, and calibration factors (c_i,UV_) were determined with the following equation:

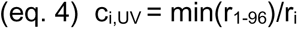

LED intensities were adjusted and measured, and this process was iterated as for the other LEDs. The final UV LED calibration factors were manually added to *optoReader.ino* at the initial set-up of the oPR.

### Correcting OD values to match Tecan plate reader

Comparison of OD from the oPR and Tecan found that, while data from both instruments were ∼linearly related, the absolute magnitudes of readings from the oPR vs Tecan varied by ∼2-fold (**Figure 5B).** To bring oPR readings in line with industry-standard Tecan readings, a transform function was generated that related oPR OD readings to expected Tecan readings. During experiments, oPR OD readings were passed through the inverse of this equation to generate final adjusted readings. The lines of best fit for the per-well data in **Figure 5B** were generated in R using the code:

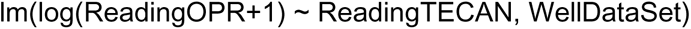

where lm() generates a line of best fit between log(ReadingOPR+1) and ReadingTECAN, and ReadingOPR represents OD readings at all concentrations for a single well. ReadingTECAN represents all Tecan readings for that same well, and WellDataSet is the data frame that these well values are stored within. A logarithmic transformation was used because it was empirically found to provide a better fit to the data, particularly at lower ODs. This formula generates the slope (m) and y-intercept (b) for that well which can then be applied to oPR OD readings (r) from the same well in future experiments using the following formula to generate TECAN adjusted results (c).

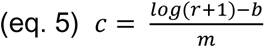

Slopes and y-intercepts can then be entered into the FeedbackFuncs.py file in order to utilize real time correction of oPR OD data for feedback enabled experiments. For experiments that do not require feedback, OD correction can be applied after completion of the experiment when processing OD outputs.

